# Ursolic Acid Inhibits Rotavirus Replication Through Modulation Of Lipid Droplet Homeostasis

**DOI:** 10.64898/2026.04.13.718265

**Authors:** Tohmé Chapini M. Julieta, Ayelén Converti, Cristian A. Pocognoni, Clara García Samartino, Benjamín Caruso, Exequiel E. Barrera, María I. Colombo, Natalia Wilke, Laura R. Delgui

## Abstract

Rotavirus (RV) replication occurs within viroplasms (VP), which are globular, membrane-less cytosolic inclusions primarily assembled by the viral NSP5 and NSP2 proteins. Among host factors, lipid droplets (LD) are strictly required for VP biogenesis. LD are ubiquitous organelles consisting of a neutral lipid core surrounded by a phospholipid monolayer and associated proteins. Ursolic acid (UA), a pentacyclic triterpenoid widely present in plants and fruits, displays multiple biological activities, including modulation of lipid metabolism, and exhibits antiviral activity against RV, as we have previously demonstrated.

Here, we investigated the molecular mechanism underlying the antiviral effect of UA. Using biophysical approaches, we first examined the impact of UA on LD formation, finding that it impairs LD biogenesis, consistent with reduced LD budding from the endoplasmic reticulum. We then employed cell-based assays to assess LD turnover and observed that UA acts as a lipolytic stimulus, leading to a marked reduction in LD abundance. Notably, we found that autophagic pathways contribute to LD degradation in the presence of UA. Finally, molecular dynamics simulations suggested that UA, owing to its intrinsic lipid-partitioning capacity, inserts into the LD phospholipid monolayer, establishing interactions with interdigitated neutral lipids.

Together, our results indicate that UA both hampers LD biogenesis and accelerates LD degradation, likely through its association with and local perturbation of the LD membrane. This dual effect leads to LD depletion, thereby impairing VP formation and ultimately inhibiting RV replication.

**IMPORTANCE:** Diarrheal diseases caused by a wide range of pathogens, including bacteria and viruses, remain a major global public health concern, accounting for more than one million deaths annually. These diseases mainly affect children under five years of age, with RV being the primary cause of diarrheal mortality in this age group. The burden is particularly severe in low- and middle-income countries.

To address RV-associated disease, coordinated public health interventions have been implemented worldwide, including improvements in water, sanitation, hygiene, the introduction of vaccines, and the widespread use of oral rehydration therapy. Although these measures have substantially reduced the global burden of diarrheal diseases, the lack of a specific antiviral treatment against RV highlights the need for continued research. The development of antiviral strategies could provide a valuable complementary approach to existing interventions and help further reduce morbidity and mortality in infected children.

## INTRODUCTION

Rotavirus (RV) is the most frequent causative agent of severe viral diarrhea and remains the leading cause of diarrhea-associated mortality in children under five years of age worldwide. Although vaccination programs have substantially reduced RV-associated morbidity and mortality, a marked disparity in vaccine efficacy persists between high-income and low- and middle-income countries (1). In 2021, RV was estimated to cause approximately 120,000 deaths among children under five globally, with the highest burden concentrated in infants between 1 and 11 months of age (2). This epidemiological landscape underscores the need to better understand the cellular and molecular mechanisms governing RV–host interactions. RV is a non-enveloped virus with an icosahedral capsid enclosing eleven segments of double-stranded RNA (dsRNA) genome (3). The error-prone nature of the viral RNA-dependent RNA polymerase, together with genome segmentation, confers a high capacity for genetic variation, facilitating the emergence of novel strains and potentially enabling vaccine escape (4). These features, combined with suboptimal vaccine performance in resource-limited settings, call for continued investigation into RV molecular biology and for the development of complementary antiviral strategies.

RV replication and assembly take place in specialized cytoplasmic inclusions known as viroplasms (VP), which are large, membrane-less, electron-dense structures. VP biogenesis critically depends on the viral non-structural proteins NSP2 and NSP5, as well as on host lipid droplets (LD). Indeed, RV infection induces a pronounced accumulation of LD that closely associates with VP (5). Pharmacological perturbation of LD metabolism disrupts VP formation and inhibits viral replication, highlighting the functional relevance of LD during RV infection (6, 7). Despite these observations, the precise mechanistic role of LD in the RV life cycle remains unclear. In this context, our group previously identified ursolic acid (UA) as a potent antiviral compound against RV, demonstrating that UA hampers the formation of VP and strongly inhibits viral replication and progeny production (7).

LD are multifunctional and highly dynamic organelles that play a central role in cellular lipid and energy metabolism (8). Structurally, they consist of a core of neutral lipids—primarily triacylglycerols and sterol esters—surrounded by a phospholipid monolayer decorated with specific proteins (9). The biogenesis of LD occurs at the endoplasmic reticulum (ER) and is driven by lipid demixing processes that are sensitive to membrane tension and phospholipid composition (10, 11). Conversely, the turnover of LD is tightly regulated by cellular metabolic demands and can be achieved through two main pathways: lipolysis, mediated by cytosolic lipases, and lipophagy, a selective form of autophagy (12–15). Importantly, chaperone-mediated autophagy (CMA) precedes both processes by selectively removing LD-coating proteins such as PLIN2 and PLIN3, thereby facilitating access of lipases and the autophagic machinery to the LD surface (16). CMA substrates containing KFERQ-like motifs are recognized by Hsc70 and delivered to the lysosomal receptor LAMP-2A for degradation. In the case of PLIN2, phosphorylation promotes its recognition by Hsc70 and removal from the LD surface, linking CMA activation to subsequent LD turnover (16–18).

Treatment of infected cells with UA results in a marked reduction in viral protein levels and a strong decrease in the production of infectious progeny. Temporal analysis revealed that UA exerts its antiviral activity predominantly during the early stages of infection, impairing VP formation and the accumulation of both structural and non-structural viral proteins, while having little effect when administered at later stages of replication (7). Given the well-established effects of UA on lipid metabolism and its amphiphilic nature, we hypothesized that its antiviral activity may stem from a direct impact on LD homeostasis, potentially through physicochemical interactions with LD membranes. In this study, we first extended our analysis by confirming the antiviral activity of UA against two additional RV strains, Rhesus RV (RRV) and bovine RV (NCDV). We then combined biophysical and cell biological approaches to elucidate its mechanism of action. Using Langmuir monolayers, we show that UA interferes with LD biogenesis by affecting their budding. Moreover, by using cell-based approaches, we observe that UA promotes autophagic pathways contributing to LD degradation. Finally, molecular dynamics simulations reveal that UA inserts into the phospholipid monolayer of LD, facilitated by the neutral lipid core, increasing the disorder of surrounding lipids. Together, our results suggest that UA disrupts LD homeostasis by impairing their biogenesis and accelerating their degradation through membrane association and local perturbation, thereby depleting LD, compromising VP formation, and RV replication.

## RESULTS

### UA exerts antiviral activity against different rotavirus strains and impairs VP formation

Given the importance of excluding cytotoxicity as a contributor to the antiviral effect, we reassessed the impact of UA on MA104 cell viability using a complementary lactate dehydrogenase-release assay, in addition to our previously reported Alamar Blue analysis (7). Treatment with 10 µM UA for 15 h did not significantly increase cytotoxicity compared with untreated or DMSO-treated cells **(Figure S1**). Having confirmed that UA was well tolerated under these experimental conditions, we next extended our previous analysis of its antiviral activity. We have previously demonstrated that UA exhibits an antiviral effect in MA104 cells infected with the simian agent 11 RV (SA-11) (7). Here we extended this analysis to two additional group A rotaviruses: the rhesus rotavirus strain RRV (G3) and the bovine rotavirus strain NCDV (G6). Treatment with 10 μM UA significantly reduced the production of infectious progeny for both strains, although the magnitude of the reduction was lower than that previously observed for SA-11 (**Figure 1A**). To determine whether this antiviral effect was also associated with impaired VP formation, we infected MA104 EGFP-NSP5 cells with SA-11, RRV, or NCDV in the presence of DMSO or 10 μM UA and quantified VP by confocal microscopy. UA treatment significantly reduced the number of VP formed by all three strains compared with the corresponding DMSO-treated controls (**Figure 1B**). Thus, despite strain-dependent differences in the reduction of infectious progeny, the inhibitory effect of UA on VP biogenesis was consistently observed across the three rotavirus strains tested, indicating that this effect is not restricted to SA-11.

**Figure 1.**
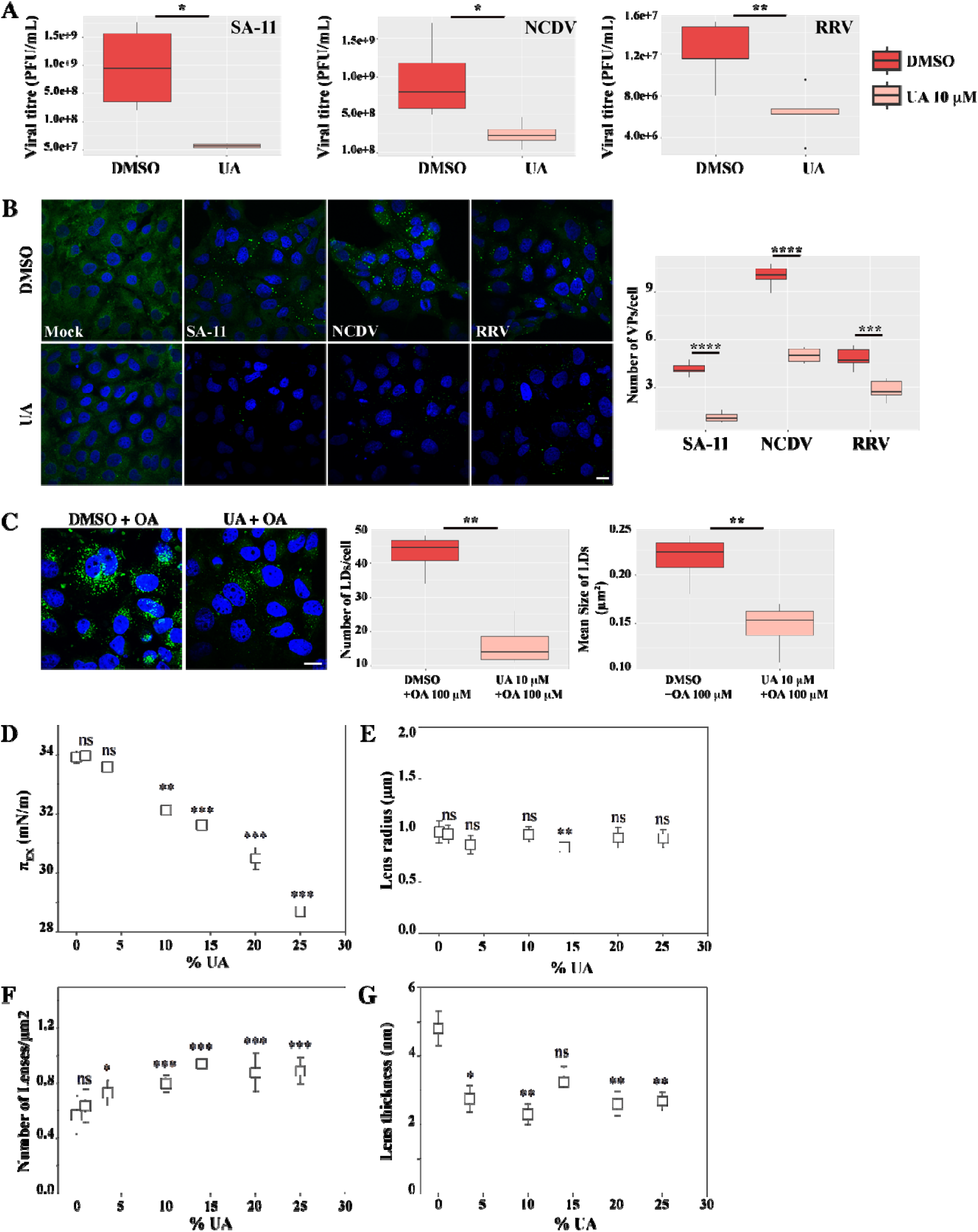
UA exerts antiviral activity against different rotavirus strains and impairs VP formation. **(A)** MA104 cells were pre-treated with DMSO or 10 μM UA for 1 h and then infected with different RV strains SA-11, NCDV and RRV at an MOI of 1 in the presence of the compounds. At 15 h post infection (p.i.), the supernatants were collected, and infectious particles were titrated using the plaque assay method. Each boxplot represents the mean titer after DMSO or 10 μM UA treatment of three independent experiments. The means were analyzed by a Student’s t-test (*p< 0.05, **p < 0.01). **(B)** MA104 EGFP-NSP5 cells were pre-treated with DMSO or 10 μM UA for 1 h and then infected with SA-11, NCDV and RRV in the presence of the compounds. At 7 h p.i., cells were fixed and analyzed by confocal laser scanning microscopy. Viroplasms were identified by EGFP-NSP5 fluorescence and quantified as the number of viroplasms per cell. Representative images are shown on the left, and boxplots on the right represent data from three independent experiments. Statistical significance between DMSO- and UA-treated cells was evaluated using Student’s t-test (***p < 0.001, ****p < 0.0001). The scale bar represents 20 μm. **(C)** MA104 cells were pre-treated with DMSO or 10 μM UA for 1 h, and then the LD accumulation was induced with 100 µM OA for 2 h. Then, the cells were incubated with BODIPY 493/503, fixed, and analyzed by CLSM. The left panel shows representative microscopy images, and the boxplots represent the mean number and size of LD of three independent experiments. The means were analyzed by a Student’s t-test (**p < 0.01). The scale bar represents 10 μm. **(D-G)** Physical characterization of lenses upon UA incorporation, obtained from compression isotherms and imaging of EPC:TG:UA monolayers **(D)** Lateral pressure at which TG are segregated from the film (π_EX_) determined at the sharp kink on the isotherms. **(E)** Lateral radius (*i.e*, from top view of the monolayer) of each individual lens. **(F)** Lens density (number of lenses/μm^2^). (**G)** Thickness of each individual lens, computed from BAM reflectivity using the equation included in Materials and Methods section. Statistical significance was evaluated for each %UA against the control of %UA=0 using the Tukey test (*p<0.05, **p<0.01, and ***p<0.001). D-G plots were generated using SigmaPlot for Windows v15 (Systat, USA).

### UA exerts a negative effect on the biogenesis of LD

Having established that UA impairs VP formation across different RV strains, we next investigated whether this effect could be linked to alterations in LD metabolism, using MA104 cells as a model. Cells were pre-treated with DMSO or 10 μM UA for 1 h and LD accumulation was subsequently induced with 100 μM oleic acid (OA). LD were labeled with BODIPY 493/503 and analyzed by confocal laser scanning microscopy. In the presence of UA, both LD number and size were significantly reduced compared with DMSO-treated cells (**Figure 1C**). This observation could result from inhibition of LD biogenesis, induction of LD degradation, or a combination of both processes.

LD biogenesis initiates with the accumulation of neutral lipids within the two leaflets of the ER membrane, forming membrane-embedded drops, from now on called “lenses”. These lenses eventually bud off from the ER, and therefore both processes, the formation and budding of the lenses, are important steps in LD biogenesis. UA is an amphiphilic molecule; thus, it likely interacts with lipid membranes, probably affecting the ability to form lenses, as well as their size and shape. To determine whether UA directly affects the physicochemical steps underlying LD biogenesis, independently of cellular signaling or other infection-associated processes, we used a reconstituted Langmuir monolayer system previously validated for studying LD biogenesis (11). This cell-free model mimics the phospholipid/neutral-lipid environment in which neutral lipids segregate within the ER membrane to form lipid lenses. Langmuir films composed of triglycerides (TG) and egg phosphatidylcholine (EPC) were compressed while monitoring surface pressure (π) and monolayer morphology by Brewster Angle Microscopy (BAM). The π at which TG exclusion occurred to form micrometer-sized lenses (π_EX_) was detected as a clear inflection point in the compression isotherm, while lenses appeared as bright structures in BAM images. A detailed description of these experiments is provided in Supplementary Text sections S1–S3.

Accordingly, control experiments were first performed. Compression isotherms of single-component monolayers (TG, UA or EPC) reproduced previously reported profiles (19, 20) and are presented in **Figure S2A**. The EPC:TG mixture was also tested, since it constitutes our model for the biogenesis of LD. Experiments for the binary mixtures containing UA (EPC-UA and TG-UA) are also shown in **Figure S2 (B, C)** and the ternary mixture EPC:TG:UA in **Figure S2D**. Representative BAM images of these films are shown in **Figure S3**. Monolayers formed with TG films exhibited lenses once π_EX_ was reached (**Figure S3 e,f**), whereas pure UA, EPC and EPC:UA mixtures monolayers did not (**Figure S3 c, d and i-l**). The films of EPC:UA collapsed at intermediate pressures (π_2_) between those of pure EPC or UA, suggesting molecular mixing of components in accordance with what was previously reported (21) (**Figure S2B and Figure S4**). On the contrary, in monolayers formed with TG:UA, lenses were formed at a π_EX_like that of pure TG films, suggesting the unmixing of the components (**Figures S2C and S4**). In summary, these experiments indicate that i) UA does not segregate into domains when mixed with EPC or TG, ii) UA does not interfere with the formation of TG lenses, and iii) UA alone does not form micron-scale aggregates or lenses that may be indistinguishable from TG lenses.

Next, we analyzed the influence of UA on the formation of lenses in the EPC:TG model. For this, monolayers composed of a ternary mixture of EPC:TG:UA were compressed. These films were homogeneous up to π_EX_, with π_EX_ progressively decreasing as the proportion of UA increased (**Figure 1D**), suggesting interactions between UA and EPC. We propose that EPC:UA interactions may lead to a sequestration of EPC by UA, promoting an enrichment of the monolayer in TG that induces a decrease in π_EX_. In this regard, a decrease in π_EX_ with the concentration of TG has been demonstrated in EPC:TG films (19, 22). To analyze the influence of UA in the shape of the TG lenses, the lateral size and the height of the lenses were quantified from the BAM images. These determinations revealed that UA did not affect their lateral size (**Figure 1E**) while inducing an increase in the number of lenses per area (**Figure 1F**). Moreover, the increase in lens density correlates with a decrease in the thickness of lenses (**Figure 1G**), indicating that a similar amount of TG molecules was expelled from the interface, though with a different distribution. These results suggest a higher tension of the membrane surrounding the lens as the percentage of UA increases (as observed from the decrease in π_EX_). The impact of UA on the shape of the lenses is explained by the fact that they form at a higher surface tension, promoting their pulling out and thus preventing them from being round. Extrapolating these results to *in vivo* conditions, they suggest that lenses in the ER membrane would be flatter in the presence of UA and therefore less prone to bud off, remaining attached to the ER. These results suggest that the presence of UA affects the budding of LD from the ER, explaining the decreased number and size of LD present in the UA-treated cells, shown in **Figure 1C**.

### UA promotes the degradation of LD

To analyze the possible effect of UA as an inducer of LD degradation, we used the following experimental scheme. We induced the accumulation of LD by the treatment with 100 µM OA for 2 h, and then the cells were incubated with DMSO or 10 µM UA. We stained the LD with the fluorescent probe BODIPY 493/503 and used CLSM to quantify the number and size of the LD. The cells treated with UA exhibited a significant reduction in both the number and size of LD in comparison to those treated with DMSO, suggesting that UA promotes the breakdown of the LD (**Figure 2A**). In an alternative approach, we assessed the fluorescent signal in these cells using flow cytometry, which reflects the extent of BODIPY 493/503 incorporation. The mean fluorescence intensity of the cells treated with UA was significantly less than that of the DMSO-treated cells (**Figure 2B**). In this experiment we incorporated cells treated only with DMSO or UA and obtained non-significant differences in the fluorescent signal, which we expected given that the basal number of LD is low in epithelial cells. Finally, to get insight into the kinetics of LD degradation promoted by UA, we incubated MA104 cells with 100 µM OA and treated with DMSO or 10 µM UA for 0, 2, 4, and 6 h. As depicted in **Figure 2C**, we observed a significant decrease in the number and size of LD in UA-treated cells from 2 h post-treatment, and this effect persisted throughout the assessment period. These observations indicate an association between the presence of UA and the promotion of LD degradation.

**Figure 2.**
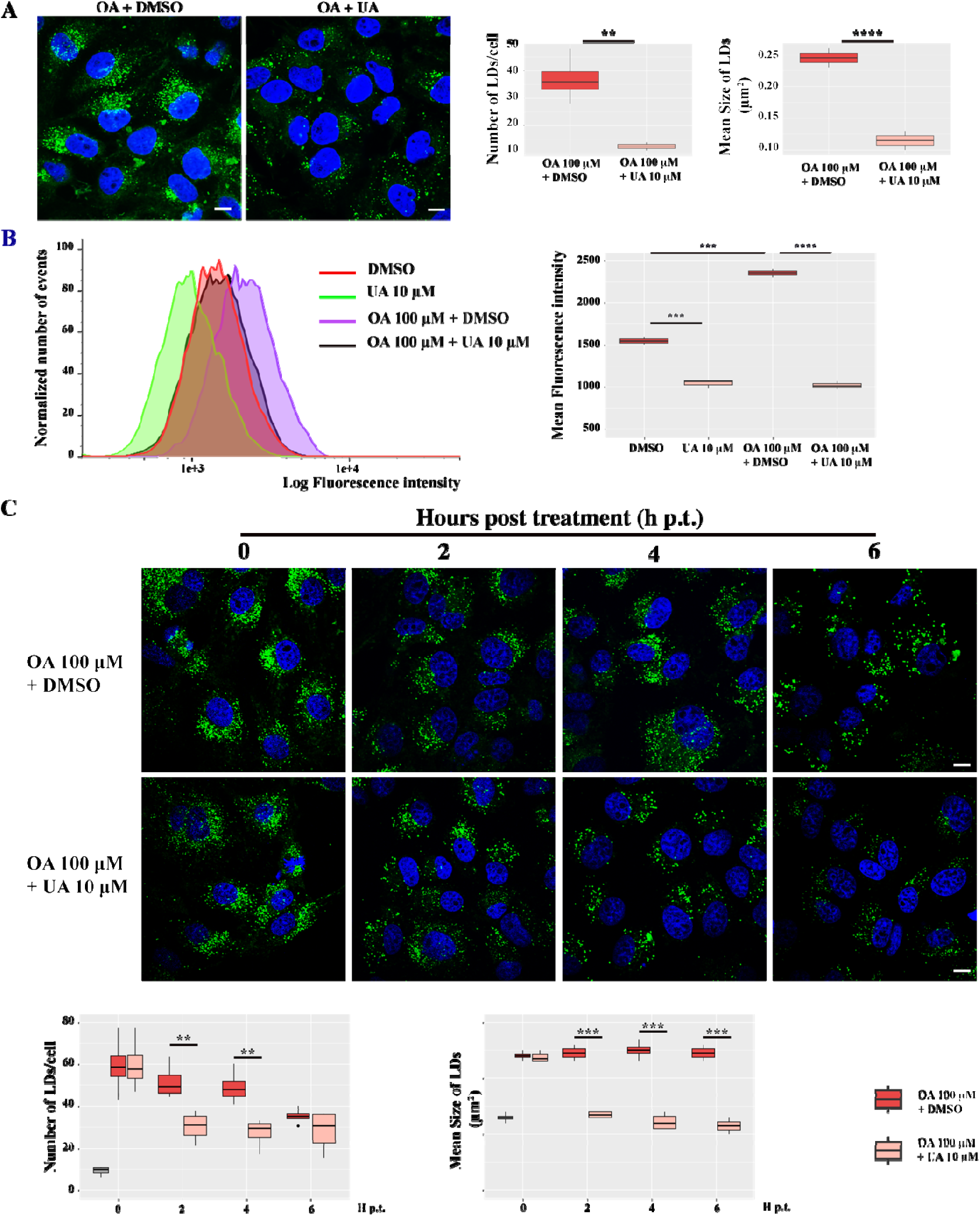
UA promotes the degradation of LD. **(A)** MA104 cells were treated with 100 µM OA for 2 h and then treated with DMSO or 10 µM UA for 1 h. Then, the cells were incubated with BODIPY 493/503, fixed, and analyzed by CLSM. The left panel shows representative microscopy images. The scale bar represents 10 µm. The boxplots represent the mean number and size of LD of three independent experiments. The means were analyzed by a Student’s t-test (**p < 0.01, ****p<0.0001). **(B)** MA104 cells were treated with 100 µM OA for 2 h and then treated with DMSO or 10 µM UA during 1 h as in (A). Additionally, cells were treated with DMSO or 10 µM UA alone. Then, they were incubated with BODIPY 493/503, and the fluorescence intensity was analyzed by flow cytometry. 10,000 events per condition were evaluated from four independent experiments. The left panel shows representative histograms, where the area under the curve corresponds to the fluorescence intensity. The boxplot represents the fluorescence intensity means for each condition. The means were analyzed by a Student’s t-test (***p<0.001; ****p< 0.0001). **(C)** MA104 cells were treated with 100 µM OA for 2 h and then incubated with DMSO or 10 µM UA during 0, 2, 4, or 6 h. Then, the cells were incubated with BODIPY 493/503, fixed, and analyzed by CLSM. The upper panel shows representative microscopy images where the scale bar represents 10 µm. The boxplots represent the mean number and size of LD of three independent experiments. The means were analyzed by a paired Student’s t-test (**p < 0.01, ***p < 0.001).

### The antiviral action of UA is mediated by its negative impact on the metabolism of LD

The LD are required for the formation of the VP (5, 23–26). Having observed that the UA interferes with the biogenesis and promotes the degradation of LD, we decided to kinetically evaluate the formation of VP in the presence of UA. To this end, we first determined the LD level throughout the early stages of the viral infectious cycle, knowledge that is currently lacking in the field. We used MA104 EGFP-NSP5 cells, a cell line constitutively overexpressing EGFP-NSP5, which is rapidly and efficiently recruited into VP upon RV infection (27). We infected the cells with RV and analyzed them at 1, 3, 5, and 7 h post infection (p.i.). We colored the LD with the red, fluorescent probe LipidTOX and used CLSM to quantify the level and size of LD. We observed a significant increase in the number and size of LD from 1 h p.i. with the peak of accumulation of LD occurring at 3 h p.i., a time point when the detection of the VP also begins. Concomitantly, we noticed that from 3 h p.i., the number of LD progressively decreased, with the lowest level observed at 7 h p.i. when the VP reached its maximum (**Figure 3**). This *biphasic* dynamic of LD during the RV life cycle, characterized by an early accumulation followed by a marked depletion as infection progresses, represents a previously unrecognized feature of the RV-LD interplay.

**Figure 3.**
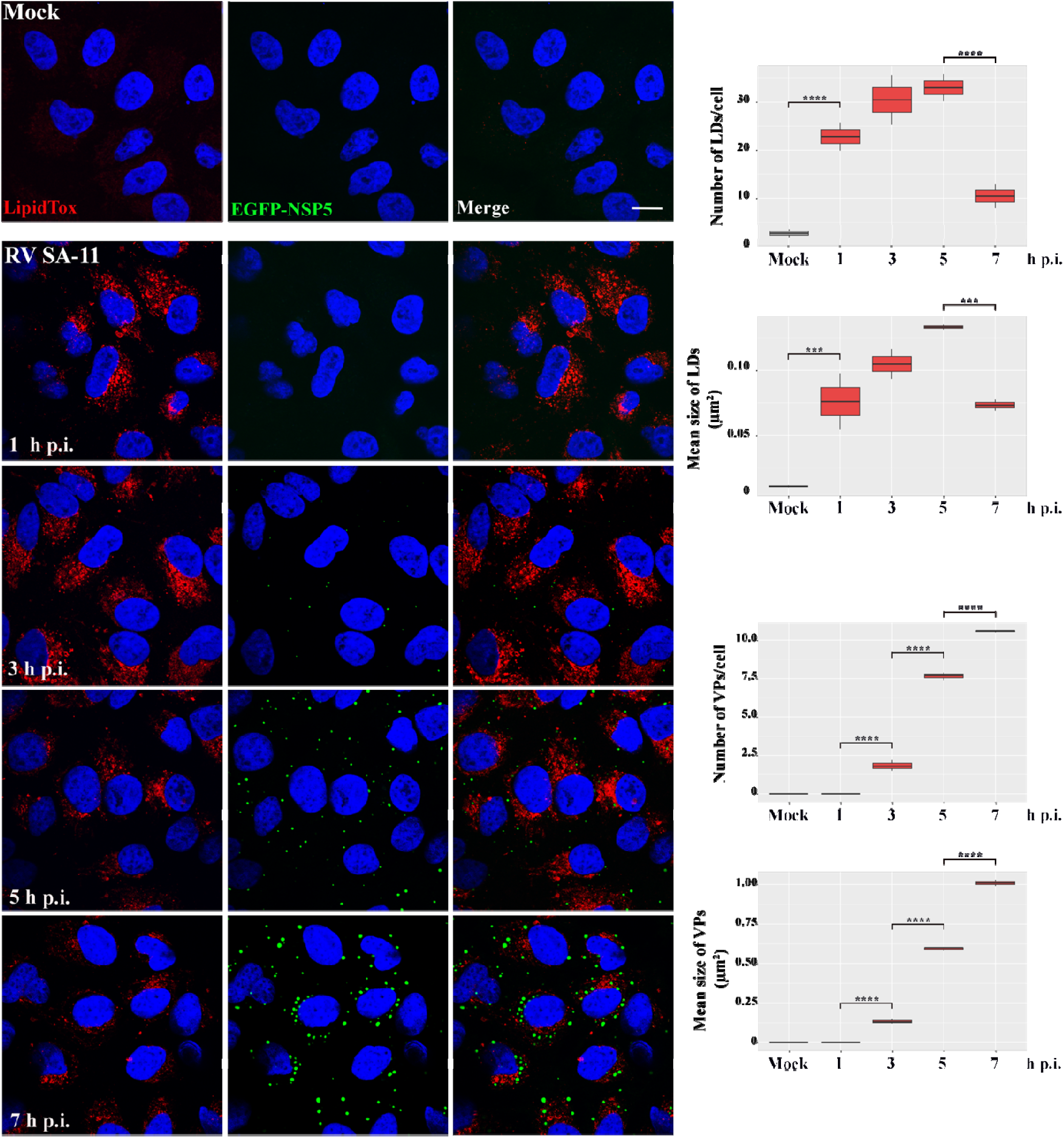
**Kinetic analysis of LD levels during the early stages of RV infection. MA104 EGFP-**NSP5 cells were infected with RV at an MOI of 1 and analyzed at 1, 3, 5, and 7 h post infection (p.i.). Then, the cells were incubated with the LD red, fluorescent probe LipidTOX, fixed, and analyzed by CLSM. The VP were evidenced by the green, EGFP-derived fluorescent dots. The left panel shows representative microscopy images where the scale bar represents 10 µm. The boxplots on the right side represent the mean number and size of LD and VP of three independent experiments. The means were analyzed by a paired Student’s t-test (***p < 0.001, ****p < 0.0001).

Then, retrieving the action of UA, we conducted a kinetic infection, where the cells were DMSO- or 10 μM UA-pretreated and then kept present during the infection. The infection was stopped at 1, 3, 5, or 7 h p.i. We stained the LD in green with BODIPY 493/503, and anti-NSP2 antibodies were used to detect the VP. We observed a significant decrease in the number and size of LD at 1, 3, and 5 h p.i., with a significant decrease in the number and size of VP in UA-versus DMSO-treated cells (**Figure 4**). To confirm our observations, we used a different approach. We pretreated and infected MA104 EGFP-NSP5 cells accordingly and used anti-PLIN2 antibodies to detect the LD. Similarly, a significant decrease in the number and size of LD at all times, with a significant decrease in the number and size of VP in UA-versus DMSO-treated cells, was observed (**Figure S5**). Overall, these observations indicate that the antiviral effect of UA results from the UA-induced reduction in the availability of LD, required for VP formation.

**Figure 4.**
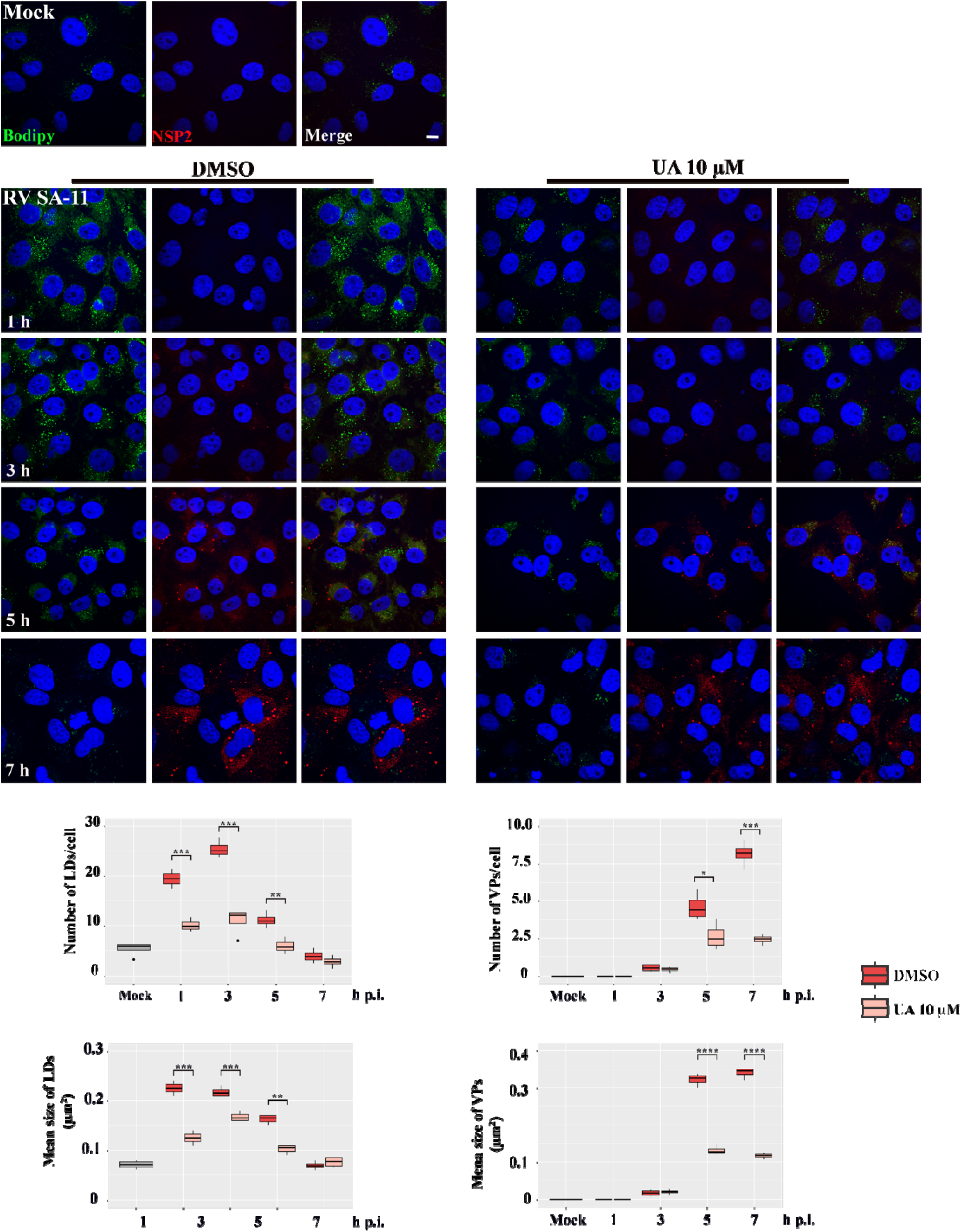
Negative impact of UA on the metabolism of LD. MA104 cells were treated with DMSO or 10 _μ_M UA for 1 h, then infected with RV at an MOI of 1 and left until 1, 3, 5, and 7 h post infection (p.i.) in the presence of DMSO (left panel) or 10 _μ_M UA (right panel). We colored the LD in green with BODIPY 493/503 and used anti-NSP2 antibodies to detect the VP. The scale bar represents 10 µm. The boxplots below represent the mean number and size of LD and VP in three independent experiments. The means were analyzed by a paired Student’s t-test (*p < 0.1, **p < 0.01, ***p < 0.001).

### CMA-related mechanisms contribute to UA-induced LD degradation

We next sought to evaluate the mechanisms involved in LD degradation induced by UA. To assess whether UA treatment was associated with increased CMA activity, we monitored the protein levels of glyceraldehyde-3-phosphate dehydrogenase (GAPDH), a well-established CMA substrate containing a KFERQ-like motif that is recognized by Hsc70 and targeted for lysosomal degradation through LAMP-2A (28). MA104 cells were treated with DMSO or 10 μM UA, and GAPDH protein levels were analyzed by Western blot at 1, 2, 4, 6, and 8 h post-treatment. UA-treated cells showed significantly lower GAPDH protein levels than DMSO-treated control at all analyzed time points. Because GAPDH is a canonical CMA substrate, this sustained reduction is compatible with increased CMA-related degradation in response to UA treatment (**Figure 5A**).

**Figure 5.**
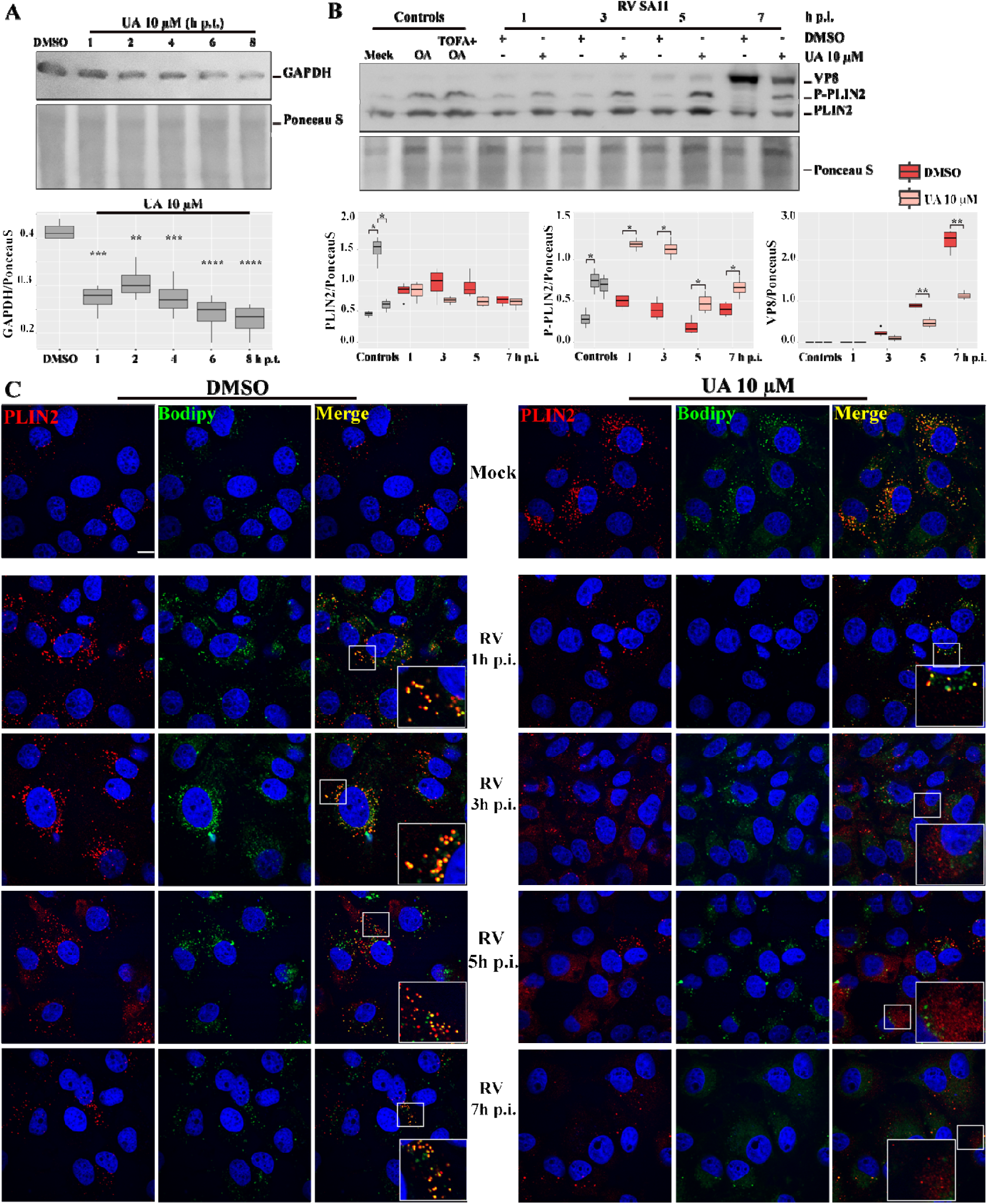
Activation of chaperone-mediated autophagy by UA. **(A)** MA104 cells were treated with 10 _μ_M UA for 1, 2, 4, 6, or 8 h. Cells were then processed for Western blot analysis, and GAPDH protein levels were detected. Ponceau S staining was used as a loading control. The boxplot was generated from four independent experiments. Statistical significance was assessed by one-way ANOVA followed by Tukey’s multiple comparisons test (**p < 0.01, ***p < 0.001, ****p < 0.0001). **(B)** MA104 cells were pre-treated with DMSO or 10 _μ_M UA and infected with RV at an MOI of 1 in the presence of the compounds. At 1, 3, 5, and 7 h post-infection (p.i.), cells were collected and processed for Western blot analysis. OA was used as a control for lipid overload and lipolytic stimulation, whereas the combination of TOFA and OA was used as a control for induction associated with inhibition of lipid synthesis. To confirm cellular infection, RV VP8 protein levels were detected. Ponceau S staining was used as a loading control. The boxplots shown were generated from four independent experiments. Statistical significance was assessed using Student’s *t*-test (*p < 0.1, **p < 0.01). **(C)** MA104 cells were pre-treated with DMSO or 10 _μ_M UA and infected with RV at an MOI of 1 in the presence of the compounds. Infection was stopped at 1, 3, 5, and 7 h post-infection (p.i.), and cells were incubated with BODIPY 493/503, fixed, and immunostained with an anti-PLIN2 antibody followed by an Alexa Fluor 555-conjugated secondary antibody. Images were acquired by CLSM. The figure shows representative images from three independent experiments. Scale bar represents 10 µm.

We next investigated whether UA modulates CMA in the context of the viral infection. To this end, MA104 cells were infected in the presence of DMSO or 10 µM UA, and p-PLIN2 levels were analyzed by Western blot at 1, 3, 5, and 7 h post-infection. OA treatment was used as a positive control, since lipid overload induced by OA increases LD number and size and promotes a lipolytic response associated with p-PLIN2 accumulation (18). As a negative control, cells were treated with 5-(tetradecyloxy)-2-furoic acid (TOFA), an inhibitor of acetyl-CoA carboxylase 1 involved in lipid biosynthesis. As expected, OA treatment resulted in a significant increase in p-PLIN2 levels compared to control conditions, whereas the OA + TOFA condition showed a modest, non-significant reduction in p-PLIN2 accumulation. RV infection was verified by detection of the viral protein VP8. Notably, RV-infected cells treated with UA showed increased p-PLIN2 signal compared with DMSO-treated infected cells, consistent with enhanced PLIN2 phosphorylation and potential targeting to CMA-related degradation (**Figure 5B**). Consistently, UA treatment exerted a clear antiviral effect during RV infection, as evidenced by a significant reduction in VP8 accumulation (**Figure 5B**).

To determine whether the UA-induced increase in p-PLIN2 levels is associated with its translocation to the cytosol, we analyzed PLIN2 subcellular distribution in infected cells treated with DMSO or UA. LD were labeled with BODIPY 493/503, and PLIN2 was detected at different times post-infection. In DMSO-treated cultures, PLIN2 largely colocalized with BODIPY 493/503-labelled LD throughout the infection time course. In contrast, UA-treated cultures showed a marked redistribution of PLIN2 away from BODIPY 493/503-positive structures and toward a more diffuse cytosolic pattern (**Figure 5C**). Because infected and uninfected cells cannot be distinguished at the single-cell level in this experiment, these observations are interpreted as qualitative evidence of altered PLIN2–LD association under UA treatment. Together with the increase in p-PLIN2 levels observed biochemically, these findings are consistent with enhanced mobilization of PLIN2 from the LD surface. Collectively, these results are consistent with the involvement of CMA-related mechanisms in UA-induced LD degradation.

### Lipophagy-mediated UA-induced degradation of LD

Lipophagy is a selective form of autophagy (12). Since UA has been reported to act as an autophagy inducer in cancer cells (29, 30), we next asked whether lipophagy could contribute to UA-induced LD degradation. To address this, we first examined the ability of UA to modulate the autophagy pathway in our cellular model (MA104 cells) using well-established autophagy modulators. Monolayers of MA104 cells were maintained under control conditions (Ctr.) or incubated for 2 h in starvation medium (Stv.), a physiological inducer of autophagy, either alone or in combination with Bafilomycin A1 (Stv. + BafA). Bafilomycin A1 blocks autophagic flux, leading to the accumulation of autophagosomes (31). Autophagosome formation was assessed by immunostaining for LC3, a well-established marker of autophagic structures, followed by CLSM. As expected, starvation induced an increase in the number of LC3-positive puncta (i.e., autophagosomes) compared with control cells, an effect that was significantly accentuated in the presence of BafA **(Figure 6A, upper panel)**. We next applied the same approach to evaluate the effect of UA. Cell monolayers were incubated for 2 h with DMSO or 10 µM UA, either alone or in combination with BafA (UA + BafA). UA treatment resulted in an increase in LC3-positive puncta, which was further significantly augmented upon co-treatment with BafA, confirming that UA acts as an autophagy inducer in MA104 cells **(Figure 6A, lower panel)**. Subsequently, we sought to evaluate the effect of UA on the autophagic pathway in the context of RV infection. To this end, we performed a kinetic infection in the presence of DMSO or 10 µM UA. During autophagy, the cytosolic form of LC3 (LC3-I) is lipidated to generate the phosphatidylethanolamine-conjugated form LC3-II, which associates with autophagosome membranes. Owing to their distinct electrophoretic mobilities, LC3-I and LC3-II can be readily resolved by SDS-PAGE, and LC3-II levels are commonly used as a proxy for autophagosome abundance. Accordingly, we performed Western blot analysis to quantify LC3-II accumulation, using starvation alone or in combination with bafilomycin A1 (BafA1) as controls for autophagy modulation. We found that cells infected in the presence of UA displayed a significant increase in LC3-II levels, compared with DMSO-treated infected cells, indicating activation of the autophagic pathway under these conditions **(Figure 6B)**. Collectively, these results suggest that UA-induced autophagy may contribute to LD degradation through lipophagy during RV infection.

**Figure 6.**
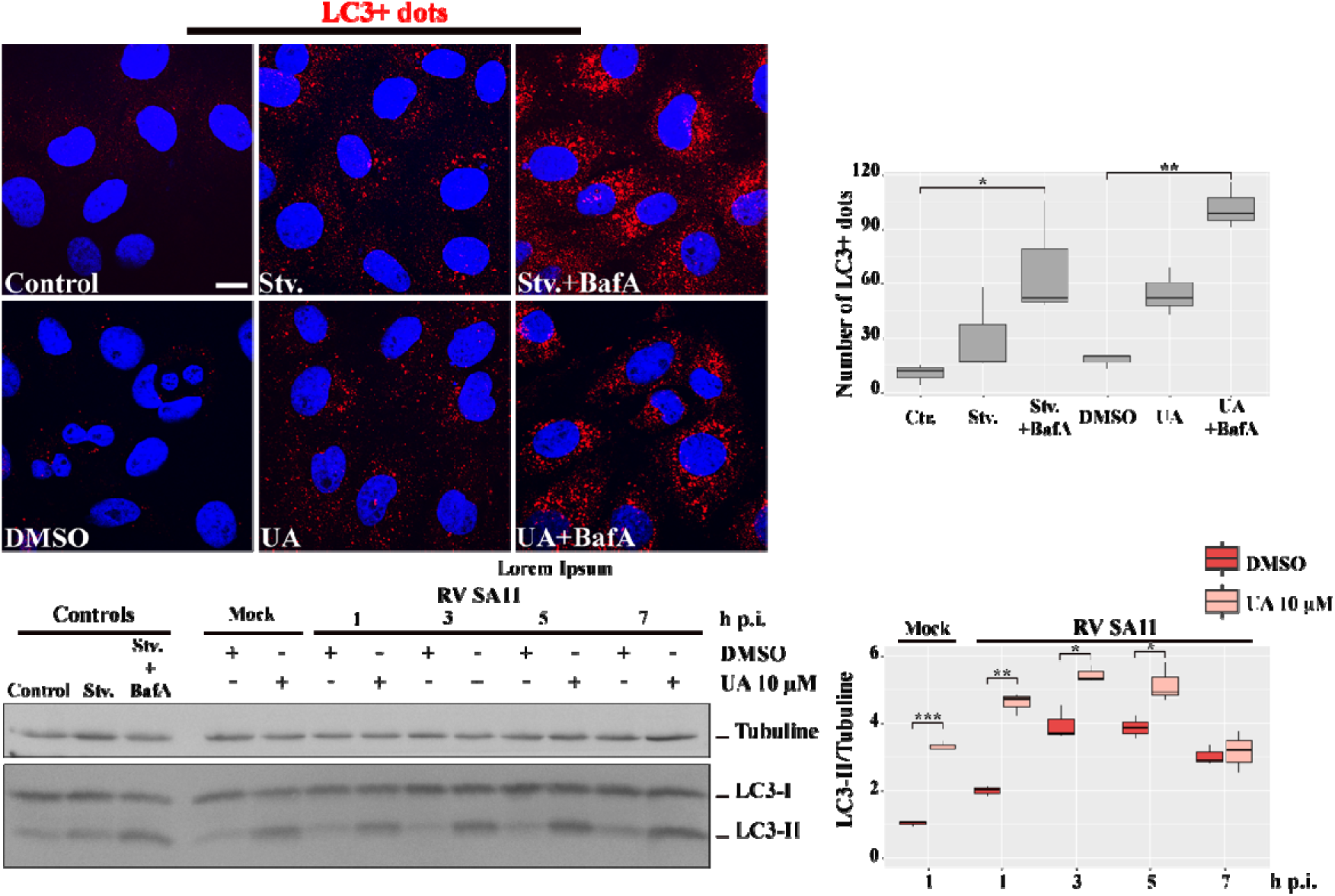
Activation of lipophagy by UA. **(A)** First row: MA104 cells were left untreated (Control), incubated for 2 h with starvation medium (Stv.), or starvation medium plus Bafilomycin A1 (Stv. + BafA1). Second row: MA104 cells were DMSO- or 10 µM UA-treated for 2 h, in the absenc or presence of Bafilomycin A1 (UA + BafA1). Cells were fixed and processed for immunofluorescence staining of LC3 (red), and nuclei were counterstained with Hoechst (blue). Representative CLSM images are shown. The number of LC3-positive puncta (LC3+ dots) per cell was quantified and is shown on the right as boxplots. Data were obtained from four independent experiments. Statistical significance was assessed using Student’s *t*-test (*P < 0.05; **P < 0.01). **(B)** Controls (first three lanes): MA104 cells were left untreated (Control), incubated for 2 h with starvation medium (Stv.), or starvation medium plus Bafilomycin A1 (Stv. +BafA1) to verify the autophagy modulation. The remaining lanes correspond to mock-treated cells or cells infected with RV strain SA-11 (MOI = 1) in the presence of DMSO or 10 µM UA. At the indicated times post-infection (1, 3, 5, and 7 h p.i.), cells were collected and processed for Western blot analysis to detect LC3-I and LC3-II. Tubulin was used as a loading control. Representative immunoblots are shown (left). All lanes shown in panel B derive from the same membrane and the same image acquisition; no lanes or blot images were spliced. Densitometric quantification of LC3-II normalized to tubulin is shown on the right as boxplots. Data correspond to four independent experiments. Statistical significance was assessed using Student’s t-test (*p < 0.05; **p < 0.01; ***p < 0.001).

### UA associates with and locally perturbs the LD monolayer

Several studies have identified molecular and biophysical properties of LD that determine their degradation route. These factors include differences in the size, which influence membrane curvature and surface tension; variations in the proteomes targeting LD subpopulations; and alterations in the lipid composition of the hydrophobic core (18, 32–39). Given its hydrophobic and amphiphilic properties, UA is expected to partition into lipid interfaces. We therefore investigated whether UA could spontaneously associate with the LD phospholipid monolayer and alter its local organization. We employed an *in silico* approach using a trilayered model of LD. This model consists of a core of neutral lipids formed of cholesterol oleate (CHYO) and trioleoylglycerol (TOG), flanked by two monolayers of 1-palmitoyl-2-oleoyl-sn-glycero-3-phosphocholine (POPC), 1,2-dioleoyl-sn-glycero-3-phosphoethanolamine (DOPE), and 1-stearoyl-2-arachidonoyl-3-glycerophosphatidylinositol (SAPI). This simplified lipid arrangement has already been employed to report different physicochemical properties of LD in similar simulations (35, 40, 41). To assess whether UA could spontaneously associate with the LD surface, four UA molecules were initially placed in the aqueous phase surrounding the LD model (**Figure 7A, panel a**). Unbiased molecular dynamics simulations were then run for 500 ns in triplicate. In two of the three replicas, UA molecules first assembled into an amphipathic tetramer before contacting the monolayer. The oligomer displayed a hydrophobic core formed by the pentacyclic triterpene scaffold and a polar surface formed by the carboxylate and hydroxyl groups exposed to the aqueous phase (**Figure 7A, panel b**). Following transient interactions with phospholipid headgroups, the UA oligomer became anchored at the LD surface through a salt bridge between the positively charged ethanolamine group of DOPE and the negatively charged carboxylate group of UA (**Figure 7A, panel b**). UA then penetrated further into the monolayer, where its position was stabilized by interactions not only with phospholipids but also with interdigitated CHYO and TOG molecules from the neutral lipid core (**Figure 7A, panel c**). At later stages of the simulation, the UA tetramer partially dissociated, and individual UA molecules remained embedded within the phospholipid monolayer (**Figure 7A, panel d**).

**Figure 7.**
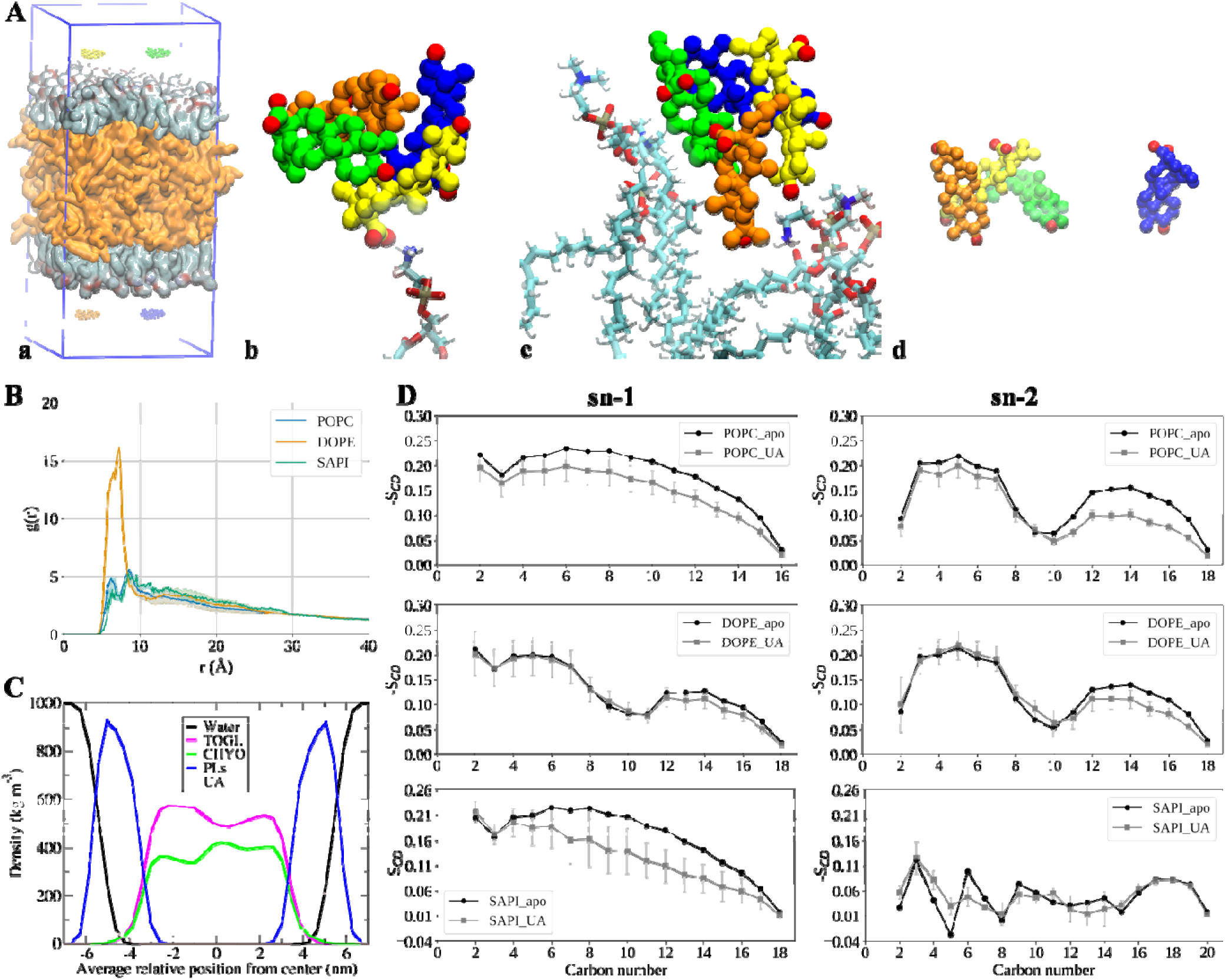
Simulated interactions between UA and a trilayered model of LD. **(A)** Graphical representation of the simulation box; phospholipids are colored in cyan, and neutral lipids in ora ge; for clarity solvent molecules are not shown **(a)**. First contact between the UA tetramer and DOPE, showing the salt bridge between the carboxylate of UA and the NH_3_^+^ **(b)**. Further insertion of UA into the monolayer, stabilized by interactions with phospholipids and interdigitated neutral lipids **(c)**. The oligomer is completely inserted into the monolayer, and one UA monomer is dissociated **(d)**. **(B)** Radial pair distribution function of the different phospholipids with respect to UA. **(C)** Average density profiles of the individual components in the LD system. UA density is multiplied by 10 for better visualization. **(D)** Acyl chain order parameters (S_CD_) of the different phospholipids’ acyl chains; a comparison with the UA-free system is shown.

The preferential interaction of UA with DOPE was quantified using the radial pair distribution function, g(r), which describes the probability of finding a given phospholipid at increasing distances from UA. DOPE showed a markedly higher probability of being located within 5–10 Å of UA, with values approximately threefold higher than those observed for POPC and SAPI (**Figure 7B**).

The time courses shown in **Figure S6A** further illustrate the interaction of UA with the LD model. Once UA molecules reached the phospholipid monolayer, they remained associated with this region throughout the simulation, where they interacted with both phospholipids and interdigitated CHYO and TOG molecules. In two replicas, UA first formed oligomers before insertion, whereas in the third replica individual UA molecules interacted directly with the monolayer. Despite these differences, UA remained stably associated with the monolayer in all three simulations.

We next examined whether UA altered the distribution of neutral lipids within the LD model. For this purpose, we calculated the density profiles of each lipid species along the z-axis, perpendicular to the phospholipid monolayers, using the last 200 ns of the three simulation replicas. In the presence of UA, the CHYO density profile became asymmetric, with CHYO molecules preferentially accumulating near the monolayer containing UA. This redistribution was accompanied by an increased TOG density at the opposite monolayer interface (**Figure 7C**). In contrast, neutral lipids remained symmetrically distributed in the UA-free control system (**Figure S7**).

To determine whether this redistribution of neutral lipids was associated with increased penetration of CHYO into the phospholipid monolayer, we quantified the degree of interdigitation between CHYO and phospholipids by calculating the overlap between their density profiles. No increase in CHYO– phospholipid interdigitation was observed in the presence of UA compared with the UA-free system (**Figure S8**).

We next evaluated whether UA insertion altered the local packing of phospholipid acyl chains. For this purpose, we calculated the deuterium order parameter (S_CD_) for POPC, DOPE, and SAPI molecules located within 7 Å of UA and compared these values with those obtained in the UA-free system. Lower values of −S_CD_ indicate reduced acyl-chain order. UA insertion produced a local decrease in lipid ordering, most clearly observed for POPC and for the sn-1 chain of SAPI, whereas the effect on DOPE was comparatively modest (**Figure 7D**). These results indicate that UA locally perturbs phospholipid packing within the LD monolayer.

To assess whether the neutral lipid core contributes to the stabilization of UA within the LD monolayer, we performed additional simulations using a phospholipid bilayer with the same composition as the LD monolayers but lacking neutral lipids. In three independent 500-ns simulations, UA showed a markedly lower tendency to remain associated with the bilayer. Most UA molecules interacted only transiently with phospholipid headgroups and subsequently returned to the aqueous phase, and stable insertion was observed only for one UA molecule in one replica (**Figure S6B**). These results indicate that the presence of the neutral lipid core favors the stable association of UA with the phospholipid monolayer. Despite the stable association of UA with the LD monolayer, the simulations did not reveal major global structural perturbations of the LD model at the UA concentration examined. The effects were mainly local, as reflected by changes in phospholipid acyl-chain order near UA. Because the Langmuir experiments showed stronger effects at higher UA proportions, simulations at higher UA concentrations may be required to capture broader membrane destabilization.

Together, the simulations support a model in which UA associates with the LD surface through interactions with phospholipid headgroups, with a preferential interaction with DOPE, and is subsequently stabilized within the monolayer by hydrophobic contacts involving interdigitated neutral lipids such as CHYO and TOG.

## DISCUSSION

In this study, by combining cell biology, biophysical methods, and molecular dynamics (MD) simulations, we show that breaking LD homeostasis led to an impaired formation of VP and RV replication. We propose that UA interferes with LD biogenesis at a stage in which neutral lipids have already accumulated within the endoplasmic reticulum membrane. Consistent with this notion, our Langmuir monolayer experiments indicate that UA alters the physicochemical properties of phospholipid interfaces in the presence of neutral lipids, supporting an interference with the budding process required for LD emergence from the ER. In agreement with these observations, MD simulations show that UA preferentially associates with phospholipid monolayers in close contact with neutral lipids, suggesting that its insertion into the cytosolic hemimembrane of the ER locally perturbs the balance of forces governing LD formation. Such perturbations would be expected not only to reduce the efficiency of LD biogenesis but also to generate LD with altered membrane properties that may affect LD size (as observed upon UA treatment on cells). These alterations may render LD more susceptible to recognition by lipolytic and autophagy-related pathways, a notion further supported by our lipophagy data, which indicates enhanced LD turnover under UA treatment. In this scenario, UA-induced defects in LD biogenesis and stability converge with enhanced degradative pathways, ultimately leading to depletion of the functional LD pool required for efficient VP formation and, consequently, for progression of the viral life cycle **(Figure 8)**.

**Figure 8.**
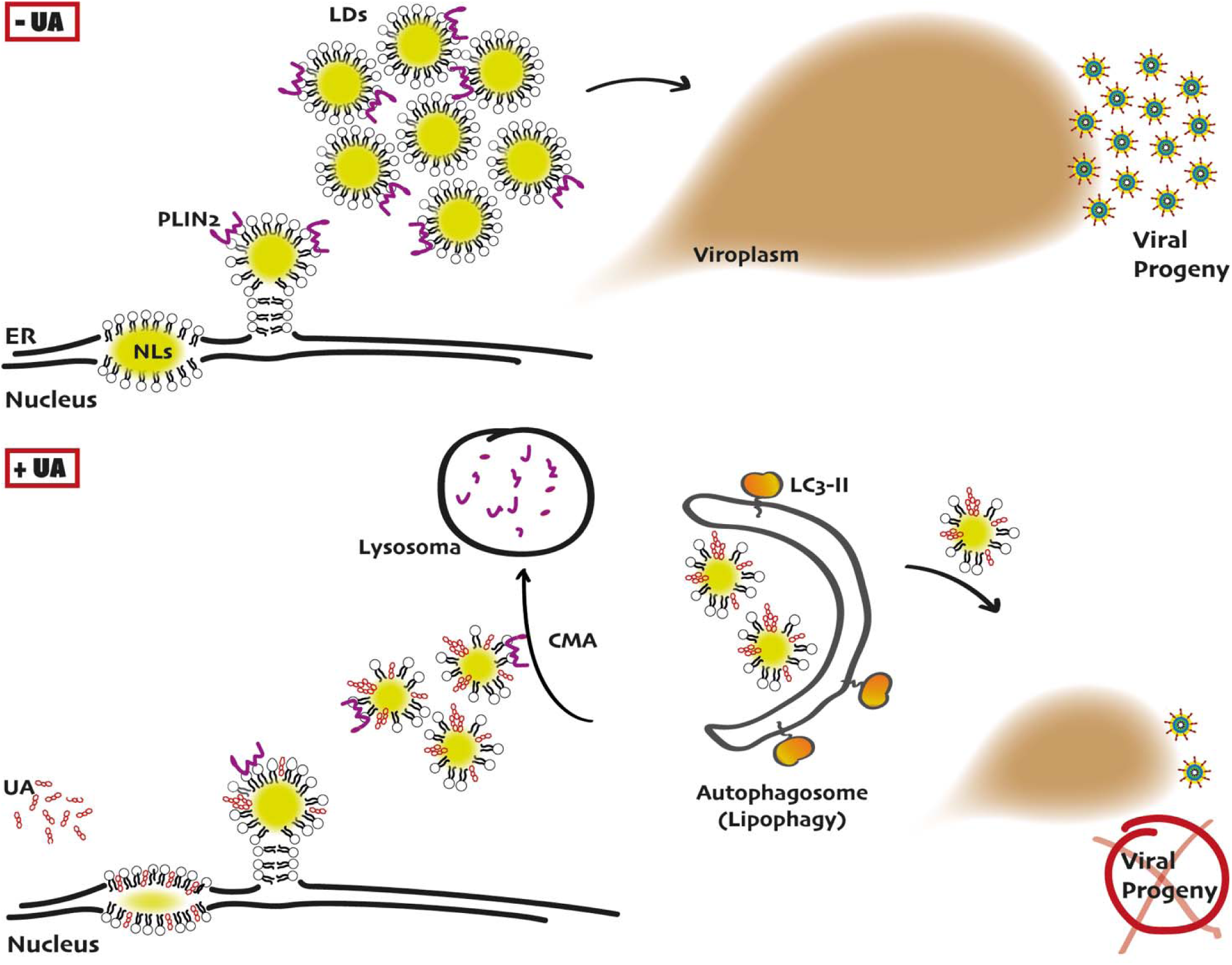
Proposed model illustrating the antiviral mechanism of UA through disruption of LD homeostasis during RV infection. Upper panel (-UA): During LD biogenesis, neutral lipids accumulate within the ER membrane, promoting the formation of the LD and the establishment of a functional LD pool that supports VP nucleation and viral replication. Lower panel (+UA): Biophysical analyses and molecular dynamics simulations show that UA preferentially associates with ER membrane regions enriched in neutral lipids. This interaction disrupts the balance needed for proper LD budding, leading to reduced LD formation. Additionally, LD that develop in the presence of UA display modified membrane characteristics and diminished size, which may favor premature degradation via lipolytic and autophagy-related mechanisms, such as chaperone-mediated autophagy (CMA) and lipophagy. The combined inhibition of LD biogenesis and acceleration of LD turnover leads to depletion of the functional LD pool below the threshold required for efficient VP formation, thereby impairing RV replication.

We characterize LD as dynamic and temporally regulated host organelles throughout the RV life cycle, revealing a previously unrecognized *biphasic* behavior marked by a pronounced early accumulation followed by gradual depletion as infection progresses. Given that epithelial cells typically contain low basal levels of LD, this initial increase is unlikely to reflect mobilization of pre-existing stores and instead suggests a virus-driven induction of LD biogenesis at early stages of infection. An intriguing question arising from these observations is whether LD actively participates in the nucleation events that give rise to VP formation. The transient accumulation of LD, together with their close spatial association, raises the possibility that LD provide a favorable physicochemical interface for the initial concentration, organization, and condensation of viral components. In this context, one may speculate that VP assembly could involve wetting-like phenomena, whereby NSP5/NSP2-rich viral condensates preferentially associate with and spread over the LD surface, thereby lowering the energetic barrier for nucleation and stabilization of replication factories, as described for phase-separated biomolecular condensates interacting with cellular interfaces (42). In this scenario, LD would function as dynamic platforms that facilitate VP assembly before being remodeled or consumed as infection advances. Future studies will be required to determine whether LD-associated membranes or lipid species directly contribute to the wetting, nucleation, and maturation of VP.

As mentioned, several studies have reported that differences in the size (concomitantly in membrane curvature and surface tension); variations in the proteomes targeting LD subpopulations; and alterations in the lipid composition of the hydrophobic core make them more prone to be degraded (18, 32–39). While our data support the notion that structurally altered LD generated in the presence of UA may be preferentially targeted for degradation via lipophagic pathways, we cannot exclude the contribution of additional signaling mechanisms converging on LD turnover. UA has been reported to modulate cellular energy-sensing pathways, including activation of AMPK, a central metabolic kinase that integrates energy status with lipid metabolism and autophagy (43). Removal of perilipins from the LD surface represents a key initiating step preceding neutral lipid degradation, and AMPK activation has been shown to induce phosphorylation of PLIN2, facilitating its association with Hsc70 and subsequent lysosomal degradation by CMA (18). Molecular docking analyses have further suggested that UA may act as a direct AMPK activator, providing an additional layer of regulation (44). In parallel, RV is known to subvert components of the autophagy machinery to support its replication. NSP4-mediated release of Ca² from the endoplasmic reticulum activates a CaMKK2–AMPK-dependent signaling pathway that promotes autophagy initiation, while inhibition of this pathway or depletion of autophagy-related proteins such as ATG3 and ATG5 reduces the production of infectious progeny (45). Subsequent studies further showed that RV replication induces LC3 lipidation and that LC3 and ATG7 contribute to efficient virus production, although this increase in LC3-II was not associated with a corresponding increase in canonical autophagosome formation (46). In addition, NSP4 exits the ER in COPII-derived vesicles that interact with LC3-II and traffic toward viroplasms, further supporting the notion that RV exploits autophagy-related membrane trafficking during its replication cycle (47). These findings, together with current models linking RV-induced LD biogenesis, viroplasm formation, and autophagy-related membrane remodeling, indicate that the interaction between RV and the autophagic machinery is functionally complex and predominantly proviral rather than simply reflecting a degradative cellular response (48).

In this context, the effect of UA is particularly intriguing. Whereas RV exploits components of the autophagy machinery to support its replication, our results suggest that UA may redirect this cellular response toward enhanced LD degradation while simultaneously impairing LD biogenesis. UA treatment increased LC3-positive structures and LC3-II accumulation, consistent with activation of autophagy-related pathways, while our biophysical data indicate that UA alters the physicochemical properties of lipid interfaces involved in LD formation. Thus, UA-mediated depletion of the LD pool likely results from the convergence of defects in LD biogenesis and membrane stability with enhanced degradative pathways, including lipophagy-and CMA-related mechanisms. In this scenario, an autophagy-related response that is normally exploited by RV may be shifted toward excessive or mistimed LD turnover, thereby limiting the availability of functional LD required for efficient VP formation and viral replication. This interpretation provides an additional mechanistic layer to the model summarized in **Figure 8**, in which impaired LD biogenesis and enhanced LD turnover jointly restrict the LD pool available for RV replication.

The antiviral mechanism described here is not restricted to a single RV strain. By extending our analyses to two additional strains of distinct origin, we demonstrate that UA-mediated disruption of LD homeostasis broadly impairs RV replication, reinforcing this mechanism. These findings underscore LD as central host factors in the viral life cycle and highlight their potential as antiviral targets. Given that LD are increasingly recognized as essential for the replication of multiple viruses, including both RNA and DNA viruses (49, 50), targeting LD biogenesis, stability, or turnover may represent a promising strategy for the development of broad-spectrum antivirals.

## MATERIALS AND METHODS

### Cell lines and viruses

MA104 Clone 1 cells (ATCC CRL-2378.1), derived from *Cercopithecus aethiops* monkey kidney epithelium, were kindly provided by Dr. Poncet (I2BC, Paris, France). MA104 EGFP-NSP5 cells were kindly provided by Dr. Burrone (ICGEB, Trieste, Italy). Cells were maintained in Dulbecco’s Modified Eagle Medium (DMEM; ThermoFisher Scientific, Waltham, MA, USA) supplemented with 10% fetal bovine serum (FBS; Internegocios S.A., Argentina, or Gibco, ThermoFisher Scientific, Waltham, MA, USA) at 37 °C in a humidified atmosphere containing 5% CO.

The simian agent 11 RV strain (SA-11) was kindly provided by Dr. Luque (UNSW, Sydney, Australia) (51). The Rhesus RV G3, as well as the bovine RV strain NCDV G6, were kindly provided by Dr.Peralta (IABIMO, Buenos Aires, Argentina). For the generation of viral stocks, RV particles were activated with 20 µg/mL trypsin (Sigma-Aldrich, Argentina) prior to infection of fully confluent MA104 cells at a multiplicity of infection (MOI) of 0.05 PFU/cell. Briefly, culture medium containing FBS was removed, cells were washed twice with phosphate-buffered saline (PBS), and viral inoculum was added for 1 h at 37°C to allow viral adsorption. After adsorption, DMEM supplemented with 1 µg/mL trypsin was added, and infection was allowed to proceed until complete cytopathic effect was observed. Viral stocks were concentrated by precipitation with 10% polyethylene glycol (PEG; Sigma-Aldrich, Argentina) and 2.3% NaCl, following the protocol described by Queiroz Fontes *et al*. (52).

### Drugs, antibodies, and reagents

Ursolic acid (UA), dimethyl sulfoxide (DMSO), 5-(tetradecyloxy)-2-furoic acid (TOFA), Bafilomycin A1 (BafA) and oleic acid (OA) were purchased from Sigma-Aldrich, Argentina. Starvation (Stv.) medium, EBSS, was purchased from Life Technologies (Argentina). All compounds were used according to the manufacturers’ instructions. Ursolic acid (UA; Sigma-Aldrich, Cat. No. U6753) was dissolved in DMSO to obtain a 47 mM stock solution. A 1.5 mM working solution was subsequently prepared in DMSO and diluted in culture medium immediately before use to obtain a final UA concentration of 10 µM. The final DMSO concentration was 0.67% (v/v), and vehicle-control cells received the corresponding concentration of DMSO. For Western blot analyses, the following primary antibodies and dilutions were used: rabbit anti-PLIN2 (1:500; ab52355, Abcam), rabbit anti-LC3 (1:1000; L7543, Sigma-Aldrich–Merck), mouse anti-GAPDH (1:5000; 60004-1-Ig, Proteintech), and rabbit anti-NSP2 and anti-VP8 (1:500), kindly provided by the group of Dr. Rodríguez and Dr. Luque (CNM, Madrid, Spain). Horseradish peroxidase (HRP)–conjugated anti-rabbit IgG and anti-mouse IgG secondary antibodies were purchased from Sigma-Aldrich (Argentina) and used at a dilution of 1:5000. All antibody dilutions for Western blotting were prepared in Tris-buffered saline (TBS) containing 0.05% Tween-20.

For immunofluorescence analysis, primary antibodies against PLIN2 and NSP2 were used at dilutions of 1:400 and 1:200, respectively. Lipid droplets were labeled using BODIPY 493/503 (D3922, Thermo Fisher Scientific, Waltham, MA, USA) or HCS LipidTOX Red (H34476, Thermo Fisher Scientific, Waltham, MA, USA) at dilutions of 1:2000 and 1:1000, respectively, and nuclei were counterstained with Hoechst 33342 (Molecular Probes, Thermo Fisher Scientific, Waltham, MA, USA) at a dilution of 1:1000. Alexa Fluor 488– and Alexa Fluor 555–conjugated secondary antibodies (Thermo Fisher Scientific, Waltham, MA, USA) were used at a dilution of 1:400. All immunofluorescence dilutions were prepared in phosphate-buffered saline (PBS).

### LDH cytotoxicity assay

Cell cytotoxicity was evaluated by measuring lactate dehydrogenase (LDH) activity released into culture supernatants using the Wiener Lab LDH-P UVI kit (Cat. #1009267), based on NADH oxidation monitored at 340 nm. MA104 cells were seeded in 24-well plates one day before treatment to reach approximately 100% confluence and were then incubated for 15 h with 10 µM UA, the corresponding DMSO vehicle concentration, or left untreated as a control condition. Maximal LDH release was induced by treatment with 0.1% Triton X-100 for 10 min at 37°C.

Following treatment, culture supernatants were collected, centrifuged for 5 min at maximum speed, and kept on ice until analysis. For the enzymatic reaction, 100 µL of Reagent A (pyruvate) was mixed with 2 µL of sample and incubated for 2 min at 37°C. Subsequently, 25 µL of Reagent B (NADH) was added and, after 30 s, 100 µL of the reaction mixture was transferred to a flat-bottom 96-well plate. Absorbance at 340 nm was recorded every minute for 3 min using a BMG LABTECH VANTAstar microplate reader. LDH activity was calculated from the rate of NADH consumption over time. Cytotoxicity was expressed relative to the maximal LDH release induced by Triton X-100.

### Confocal immunofluorescence microscopy

MA104 or MA104 EGFP-NSP5 cells were seeded onto 13-mm glass coverslips. Upon reaching confluence, cells were treated and infected according to the corresponding experimental protocols. Cells were then fixed with 4% paraformaldehyde (PFA) for 20 min at room temperature, washed twice with phosphate-buffered saline (PBS), and subsequently blocked and permeabilized using a bovine serum albumin (BSA)/saponin solution for 20 min at room temperature. When BODIPY 493/503 or LipidTOX was used to label lipid droplets, staining was performed prior to fixation. Cells were incubated overnight at 4°C in a humidified chamber with the appropriate primary antibodies. After five washes with PBS, cells were incubated with the corresponding secondary antibodies for 1.5 h at room temperature in a humidified chamber. Following five additional washes with PBS, coverslips were mounted using a Mowiol mounting medium containing Hoechst 33342 (1 µg/mL).

### SDS-PAGE and Western blot analysis

Cells were harvested in phosphate-buffered saline (PBS) using cell culture scrapers and collected by centrifugation at 3,000 rpm for 5 min. Cell pellets were resuspended in a Laemmli sample buffer and boiled at 100°C for 10 min. Proteins were separated by SDS-polyacrylamide gel electrophoresis (SDS-PAGE) using 10% polyacrylamide gels and subsequently transferred onto nitrocellulose membranes (Thermo Fisher Scientific, Waltham, MA, USA). Membranes were blocked with 5% non-fat dry milk in Tris-buffered saline containing 0.05% Tween-20 (TBS-T) for 30 min at room temperature and incubated overnight at 4°C with the appropriate primary antibodies diluted in blocking buffer. After three washes of 5 min each with TBS-T (0.05%), membranes were incubated with the corresponding horseradish peroxidase (HRP)-conjugated secondary antibodies for 1.5 h at room temperature.

Immunoreactive bands were detected using an enhanced chemiluminescence (ECL) substrate (Pierce ECL Western Blotting Substrate; Thermo Fisher Scientific, Waltham, MA, USA) and visualized with a LAS-4000 imaging system (Fujifilm). Band intensities were quantified using ImageJ software (MacBiophotonics version) (53).

### Virus titration by plaque assay

MA104 cells (2.5 × 10□ cells/well) were seeded in 24-well plates in DMEM supplemented with 10% fetal bovine serum (FBS) and grown to confluence within 24 h. Monolayers were washed twice with PBS and infected with serial dilutions of RV solutions. Viral stocks were activated with trypsin (20 µg/mL) for 30 min at 37°C and subsequently serially diluted (1:10) in DMEM prior to infection. Viral adsorption was allowed to proceed for 1 h at 37°C. After adsorption, cells were overlaid with a mixture of 50% 2× DMEM and 50% low-melting-point agarose (1.4%; Invitrogen, Thermo Fisher Scientific, Waltham, MA, USA) supplemented with trypsin (1 µg/mL). Plates were incubated at 37°C for 7 days. Monolayers were then fixed with 10% formaldehyde for 1 h at room temperature and stained with crystal violet for 15 min to visualize plaques. Viral titers were calculated and expressed as plaque-forming units per milliliter (PFU/mL). For statistical analyses, viral titers were normalized to the control condition, which was set to 100%.

### Flow cytometry analysis

MA104 cells (8 × 10□ cells/well) were seeded in 6-well plates and grown to confluence. Cells were washed twice with PBS and incubated with DMEM containing OA (100 µM) for 2 h to induce LD accumulation. Control cells were treated with DMSO. Subsequently, OA was removed, and cells were incubated with DMEM containing either DMSO or UA (10 µM) for 1 h. Cells were then stained with BODIPY 493/503 for 15 min at 37°C, harvested by scraping in PBS, and fixed in suspension with 4% paraformaldehyde for 20 min at room temperature. After fixation, cells were centrifuged at 3,000 rpm for 5 min, resuspended in PBS, and adjusted to a final volume of 300 µL. A total of 10,000 events per condition were acquired using a BD FACSAria III flow cytometer (Becton Dickinson). Data acquisition was performed using FACSDiva software, and data analysis was carried out with FlowJo software.

### Antiviral Assay

Confluent MA104 cells were pre-treated with 10 μM UA or DMSO for 1 h before infection with trypsin-activated RV at a MOI of 1 PFU/cell. After adsorption, viral inocula were removed, and cells were maintained in culture media containing 10 μM UA or DMSO, supplemented with 1 μg/mL trypsin. At 15 h p.i., the supernatants were collected to perform extracellular virus titration as described in this section.

### Statistical analysis and graphing

All boxplots and statistical analyses were performed using RStudio software (54), with the ggplot2 (55) and rstatix (56) packages. Details of the statistical tests used are provided in the corresponding figure legends.

### Model system for LD biogenesis

Two types of lipids were used: triglycerides (TG) and L-α-phosphatidylcholine from egg yolk (EPC). TG were obtained from the purification of walnut oil as described previously (19). The fatty acid composition has been previously characterized, together with its surface physical properties (19), which present almost identical properties to other liquid pure triglycerides (e.g. triolein). EPC was purchased from Avanti Polar Lipids (Alabaster, Alabama). Solvents used were of analytical grade.

### Langmuir films for compression experiments and BAM imaging

Monomolecular layers were prepared and monitored as described previously (19) using a Minitrough II (KSV Instruments Ltd., Finland) equipped with Teflon™ trough, Delrin barriers, and platinum Wilhelmy plate as surface pressure sensor. Between 10 and 30 µL of an organic (chloroform: methanol 3:1) solution of lipids was spread over a water subphase. About 5 min were allowed for the solvent evaporation before starting the compression. Films were compressed (or decompressed in some specified cases) isometrically at a constant rate of 10 mm/min. Reproducibility was ±3 Å and ±0.2 mN/m for molecular area and π, respectively. Before each experiment, the Langmuir trough was rinsed and wiped with 96% ethanol and several times with ultrapure water (18 MΩ, with the Osmoion system by Apema, Argentina). The absence of surface-active compounds in the pure solvents and in the subphase solution was checked before each run by reducing the available surface area to less than 10% of its original value after enough time was allowed for the adsorption of possible impurities that might have been present in trace amounts.

All the films were continuously observed by Brewster Angle Microscopy (BAM) with an EP3 Imaging Ellipsometer (Accurion, Goettingen, Germany) and a 20X objective (Nikon, NA 0.35). Images were acquired with the BAM settings configured to quantify the surface reflectivity. For this, the analyzer and polarizer were set to 0° relative to the plane of incidence, and the laser shutter and signal gain were adjusted to achieve non-saturated images with structures presenting well-defined edges. For this setup, a system calibration was performed at the clean interface before each experiment. Experiments were performed at 23 ± 1°C.

For characterization of lateral radius and number of lenses/µm^2^, at least three full (171680 µm^2^) images were analyzed for each of three replicas per monolayer composition. Fiji (ImageJ) was used to binarize (thresholding) images and automatically detect and measure lenses, while ensuring accuracy against the raw images.

### Thickness (*t*) of TG lenses at the air-water interface determination

The thickness (t) calculation was based on the procedure previously described (11, 57, 58), from the reflectivity value (Rp) of the incident light (λ=530 nm) over the center of a single collapsed structure (CS), according to the above equation, where η, η1 and η2 are the refractive indices of CS, air and subphase, respectively, and θB is the water Brewster angle (experimentally determined in each experiment):

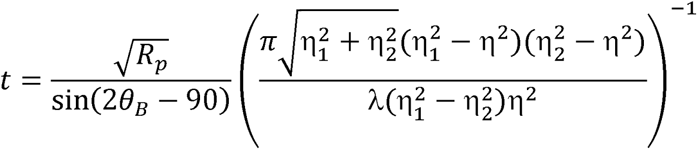

Refractive indexes are η1=1 and η2 =1.33 and η=1.47. The latter was assumed as the refractive index of bulk TG (11, 59), which was determined experimentally by assessing the Brewster angle of an air/TG interface, using a small volume black trough (60). The refractive index showed a high reproducibility (11). The gray level at the center of lenses randomly distributed in each BAM image was determined. At least three images were acquired and analyzed at each π, in three independent films.

### Trilayered-model system setup

The LD model was built with PACKMOL (61) replicating the composition proposed by Braun et *al.* (41). This system consisted of a central core of neutral lipids composed of 220 CHYO and 220 TOG molecules, located between two phospholipid monolayers each formed by 88 POPC, 37 DOPE, and 10 SAPI molecules. To improve the packing of CHYO and TOG, the neutral lipid core was surrounded by water and equilibrated via molecular dynamics simulations until stable density values. Similarly, the monolayers were first built as a bilayer patch, surrounded by water, and simulated until reaching equilibrium area per lipid values. After these first MD steps, each monolayer was separated and used as building blocks with the neutral lipid core to build the simplified trilayered LD model.

### Molecular dynamics experiments

Molecular dynamics simulations were performed using the GROMACS 2023.1 engine (62) and the Charmm36 force field (63). We employed parameters for CHYO and TOG previously published (64, 65). Parameters for UA were generated using the CHARMM General Force Field (CGenFF) via the CGenFF software (66). Systems were solvated with TIP3-P water molecules, adding potassium and chloride counterions to reach ionic concentrations of 150 mM. To avoid steric clashes systems were energy minimized using 5000 iterations of the steepest descent algorithm. Then, the system’s pressure and temperature were stabilized at 1 atm and 303 K using the C-rescale barostat and the V-rescale thermostat, respectively. The selected pressure coupling type was semi-isotropic. Simulations were run in triplicates for 500 ns using an NPT ensemble. For the ligand systems, 4 molecules of UA were placed in solution, divided into two pairs with distances of 2 nm from the top and bottom monolayers. UA were distanced by 4 nm between their geometric centers.

Visualization and rendering of images were done with VMD (67). Analysis of the structural properties of the LD model was performed over the last 200 ns of simulations. We selected this time window because in all replicas UA inserted into the monolayers after that simulated period. The radial distribution function was calculated with the g(r) GUI plugin, included in VMD. For this, we selected the carboxylate’s C atom from UA and the P atom from each phospholipid. Deuterium order parameters were calculated using the calc_op TCL script (https://www.ks.uiuc.edu/Research/vmd/mailing_list/vmd-l/att-14731/calc_op.tcl). Density profiles were calculated employing the GROMACS analysis tool gmx density. The degree of interdigitation (λ_id_) between CHYO and phospholipids was calculated by measuring the area of the intersecting density profiles of each species. To do this an overlap parameter (ρ_ov_) is first calculated as:

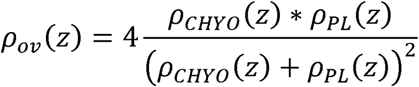

Then, ρ_ov_ is integrated along the z-axis over the simulation box to obtain λ_id_:

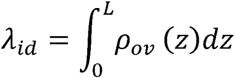

## ACKNOWLEDGMENTS AND FUNDINGS

This project was partially supported by the National Agency for Scientific and Technological Promotion, Ministry of Science, Technology and Innovation (MinCyT), through grants PICT 2016-0528 and 2019-01324 to L.R.D; the National Scientific and Technical Research Council through grants PIP 2015-2017 11220150100114CO and 2021-2023 11220200103139CO to L.R.D.; and the National University of Cuyo through grants 2016-2018 M029, 2019-2021 M071 and 2022-2024 M012 to L.R.D. The BAM experiments were performed at CEMINCO (CENTRO DE MICRO y NANOSCOPIA DE CORDOBA, Argentina), integrated into the Microscopy National System of the MinCyT. We used computational resources from CCAD—Universidad Nacional de Cordoba (https://ccad.unc.edu.ar/), which are part of SNCAD—MinCyT, República Argentina. At the IHEM, we sincerely appreciate Elisa Bocanegra, Norberto Domizio, and Jorge Ibañez for valuable technical assistance in CLSM handling.

## DATA AVAILABILITY

All data supporting the findings of this study are included in the article and its Supplementary Material.

