## Supplementary Figures for "Ursolic Acid Inhibits Rotavirus Replication Through Modulation Of Lipid Droplet Homeostasis"

**Figure S1**

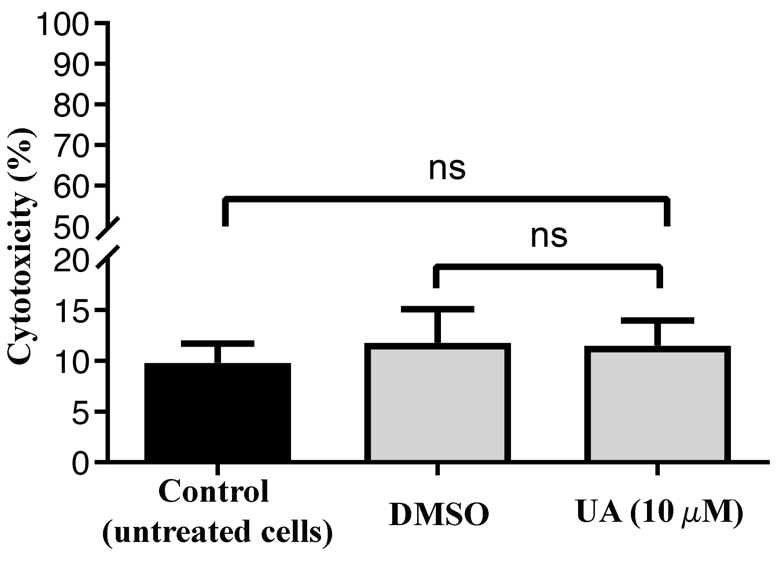

**Figure S1. Cytotoxicity analysis of MA104 cells following UA treatment.** MA104 cells were left untreated or treated with DMSO vehicle or 10 µM UA for 15 h. Cell cytotoxicity was assessed by measuring LDH release into culture supernatants through NADH oxidation at 340 nm, as described in Materials and Methods. Data are expressed as percentage of cytotoxicity and represent mean ± SD from three independent experiments. Statistical analysis was performed by one-way ANOVA followed by Tukey’s multiple-comparisons test. No significant differences were observed among untreated, DMSO-treated, and UA-treated cells.

**Figure S2**

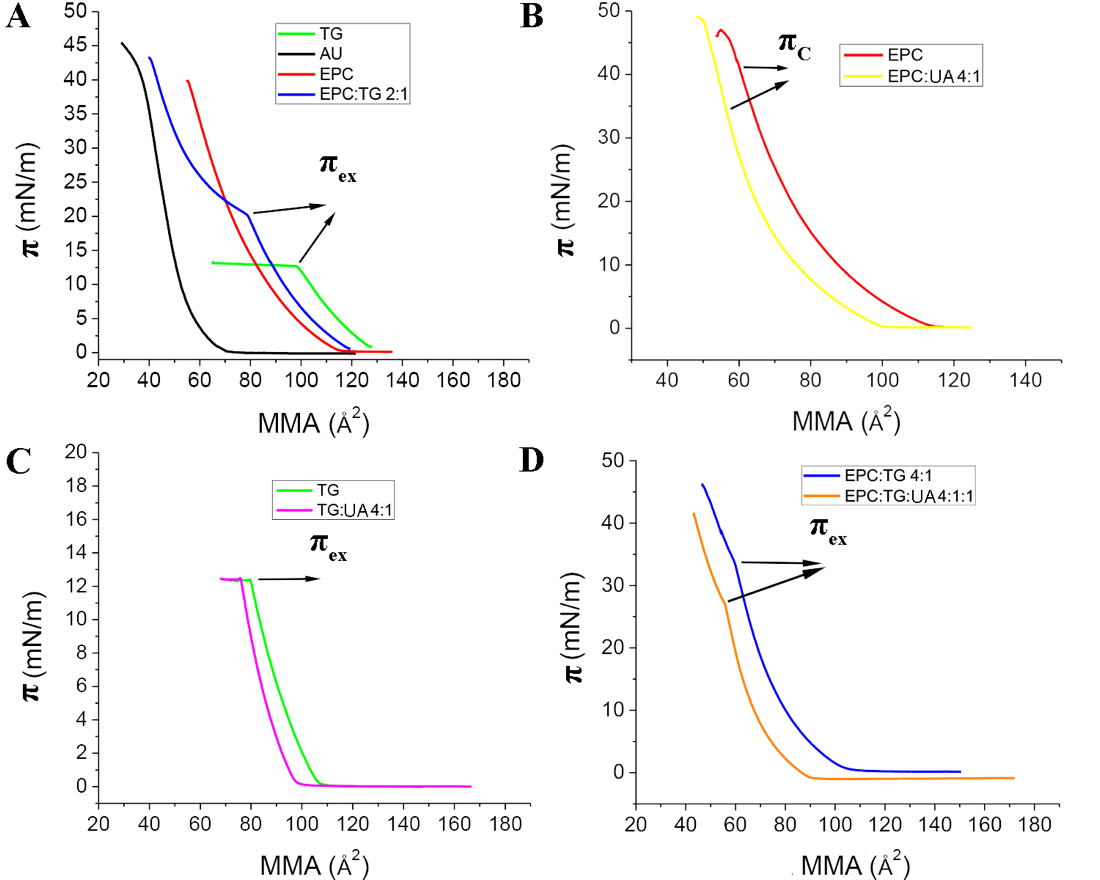

**Figure S2. Compression isotherms of monolayers. A)** Isotherms for individual components and the binary mixture of EPC:TG. Kinks on the isotherms correspond to the point $\pi_{EX}$ where TG is excluded from the monolayer into liquid lenses floating at the interface. **B, C, D)** Effect of the incorporation of UA in the lipid spreading solution. UA is mixed with EPC (**B)**, TG **(C)** or the binary mixture EPC:TG **(D).** On panel B, $\pi_{C}$ is determined as the pressure at maximum in the compressibility modulus $K=-\left( \frac{\partial\pi}{\partial MMA} \right)_{T}MMA$. Plots were generated using OriginPro v2024 (OriginLab, USA).

**Figure S3**

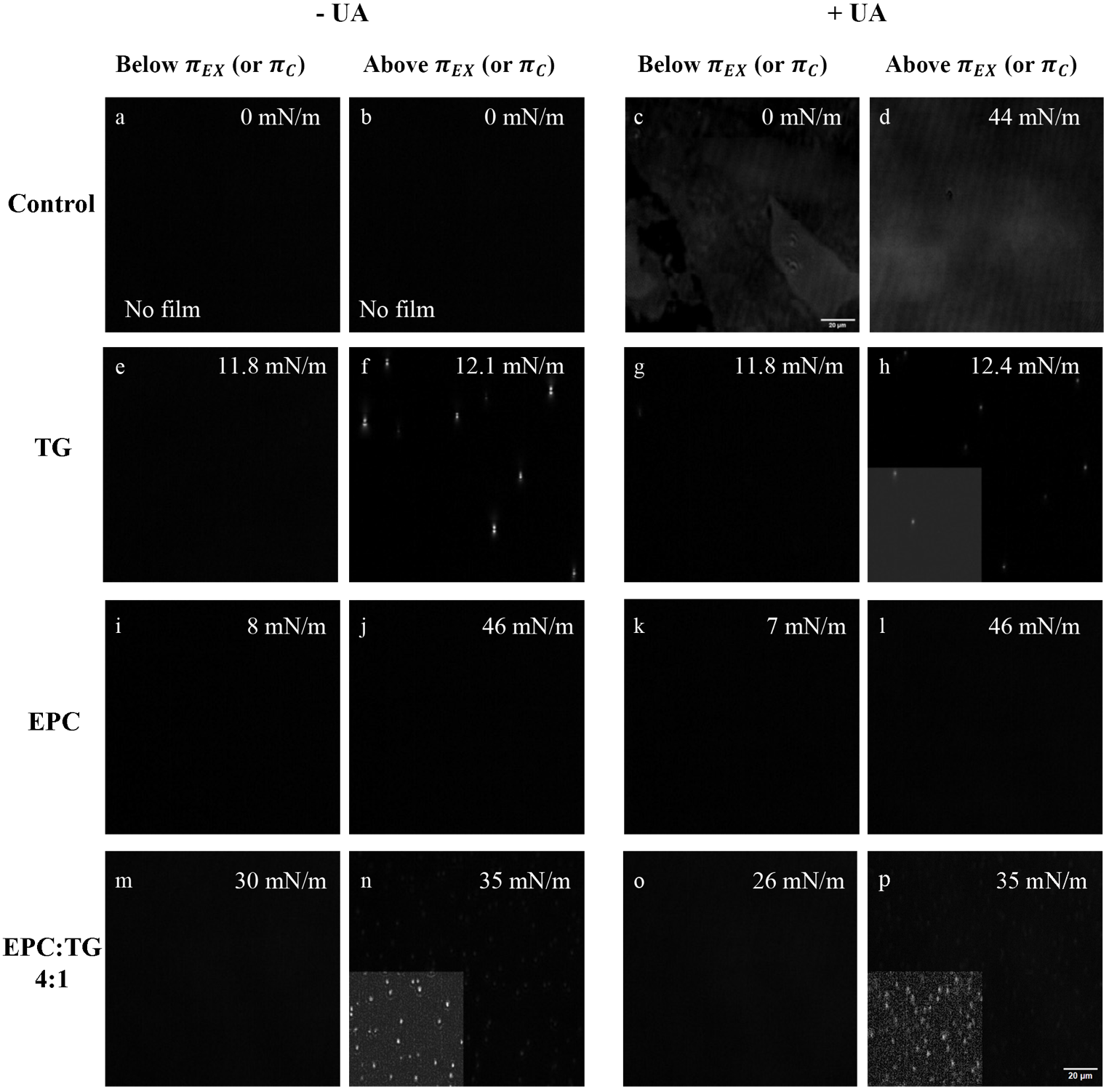

**Figure S3. BAM images of monolayers.** Different compositions (each row), with (+UA) or without (-UA) are shown. Images were acquired at lateral pressures below or above $\pi_{EX}$ (when TG were present) or $\pi_{C}$ (in the absence of TG). Each lateral pressure is indicated on each image. Scale bar corresponds to all images (20 μm). The insets in panels h, n, and p show a part of the image with adjustments in brightness and contrast for facilitating the observation of lenses (using ImageJ software). **Control Row:** images showing that spreading of the solvent (chloroform: methanol 3:1) onto the surface led to neither film formation (no rise in lateral pressure after compression) nor bright spots on the images **(a,b)**. Control of UA at 0 and 44 mN/m, at 70 Å and above monolayer collapse, respectively **(c,d)**. **TG Row:** images of TG monolayers alone or with UA. Note that UA only interferes by lowering the brightness of the lenses **(e-h)**. **EPC Row:** images of EPC monolayers alone or with UA. Note that EPC:UA does not show micron-scale structures at high pressures, as in EPC:TG. **EPC:TG** **Row:** images of the model (EPC:TG) for studying lens biogenesis in the absence and presence of UA. A clear-cut difference between monolayers below **(m)** and above **(n)** $\pi_{EX}$=34 mN/m, where lenses appear, can be observed. This behavior was conserved in the presence of UA but at lower pressures (28 mN/m) and with less bright lenses.

**Figure S4**

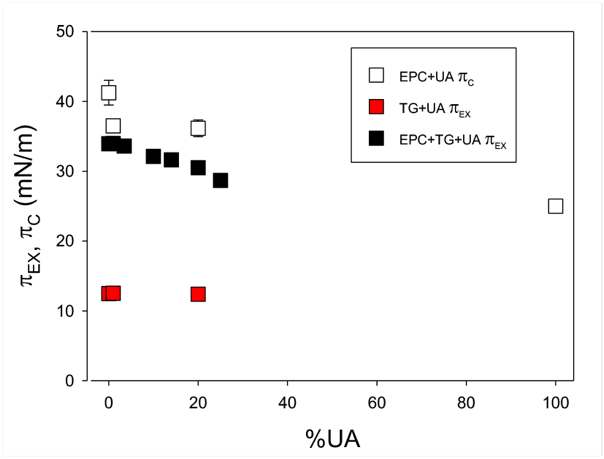

**Figure S4. Effect of UA on transition pressures of all the systems.** π_C_ corresponds to the collapse of the monolayers and is used for systems without TG, whereas π_EX_ is used for TG-containing systems and is associated with the exclusion of TG molecules into liquid lenses (observed as bright spots) while a monolayer persists at the interface. The depicted transition surface pressures correspond to monolayers composed of TG, TG:UA, and the first transition of EPC:TG:UA (the latter points being the ones plotted in **Figure 1D** and included here for comparison as black squares). Plot generated using SigmaPlot for Windows v15 (Systat, USA).

**Figure S5**

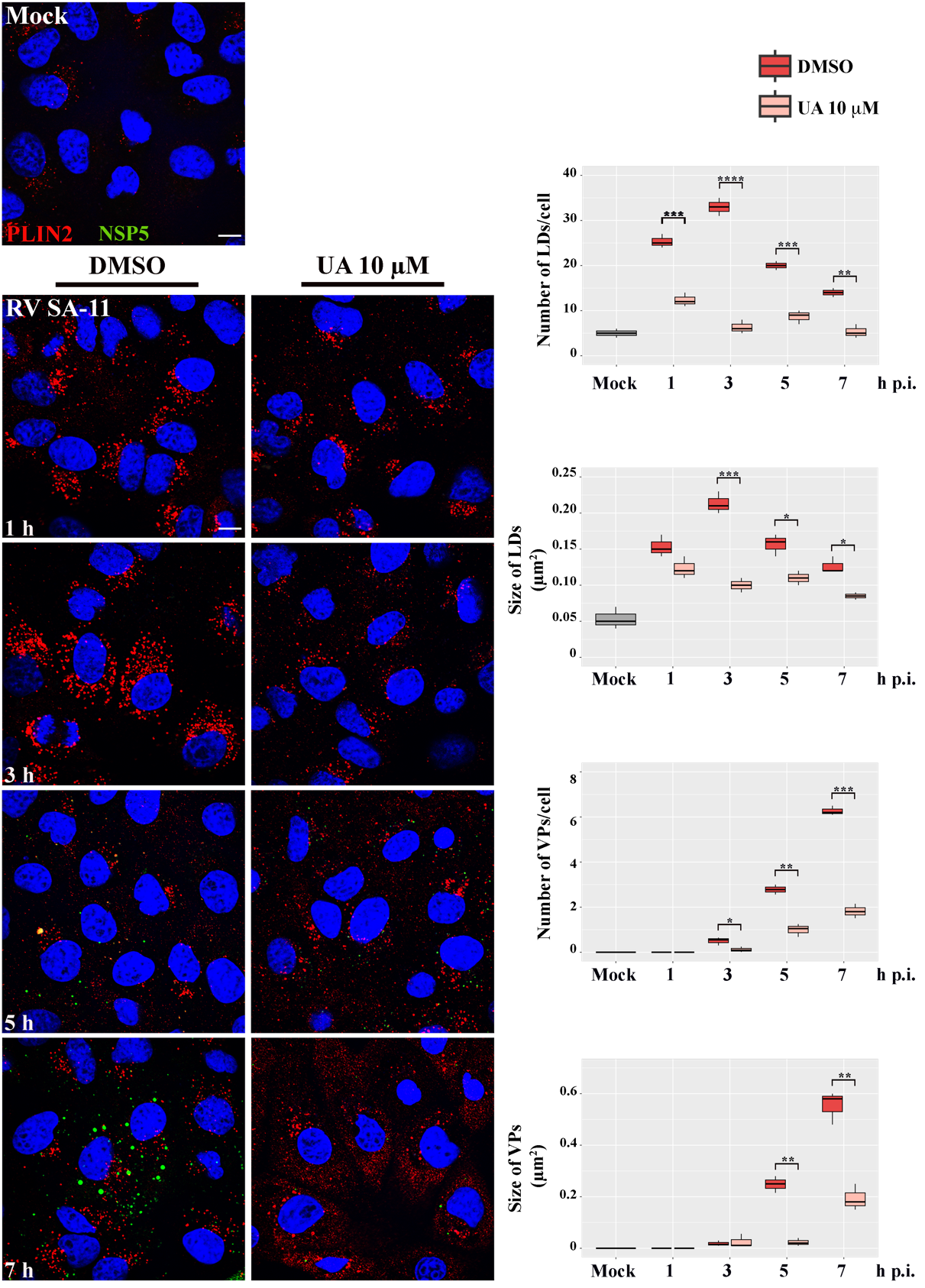

**Figure S5. Negative impact of UA on the metabolism of LD.** MA104 cells were treated with DMSO or UA 10 μM during 1 h, then infected with RV at an MOI of 1 and left until 1, 3, 5, and 7 h post infection (p.i.), in the presence of DMSO or 10 μM UA. We used anti-PLIN2 antibodies to detect the LD (red). The VP were evidenced by the green, EGFP-derived fluorescent signal. The panel shows representative microscopy images where the scale bar represents 10 µm. The boxplots represent the mean number and size of LD and VP of three independent experiments. The means were analyzed by a paired Student’s t-test (*p < 0.1, **p < 0.01, ***p < 0.001, ****p<0.0001).

**Figure S6**

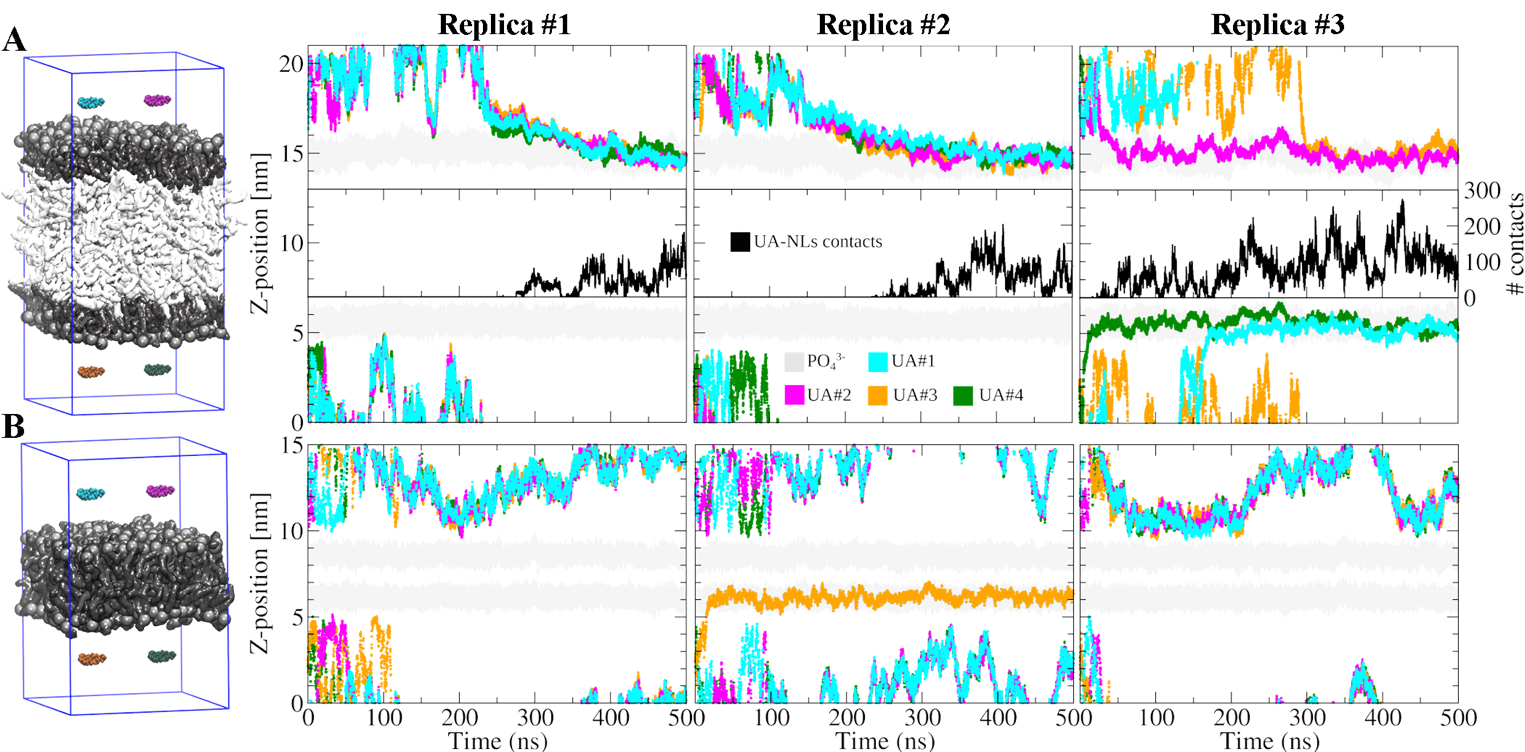

**Figure S6. Timelines showing the z-position of UA within the simulation box.** The plots illustrate the proximity of four UA molecules to phospholipids (PLs) in a lipid droplet (LD) model **(A)** and in a bilayer model **(B)**. Each column corresponds to an individual simulation replica. In panel (A), intermolecular contacts between UA and neutral lipids (NLs) are shown in the center of each panel.

**Figure S7**

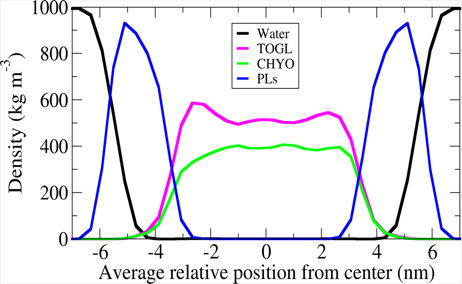

**Figure S7.** Average density profiles of the individual components in the apo LD control system.

**Figure S8**

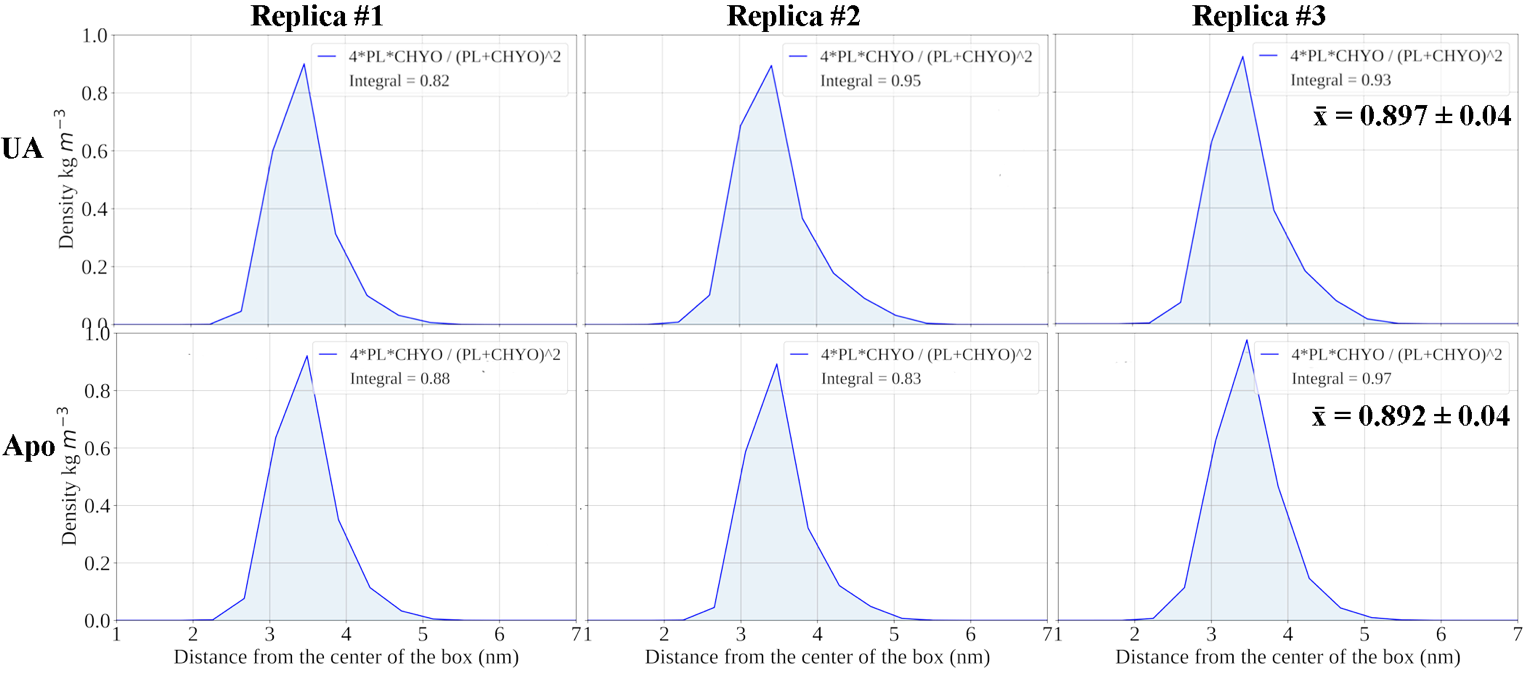

**Figure S8. Interdigitation density profiles between CHYO and phospholipid molecules.** Data are shown for each simulated replica in both UA-bound and apo systems. Mean values ± SEM are displayed in the right panels.
