## Supplementary Text for "Ursolic Acid Inhibits Rotavirus Replication Through Modulation Of Lipid Droplet Homeostasis"

**S1. Langmuir films as a model for LD biogenesis.** It is accepted that lipid droplet (LD) biogenesis in the ER is driven by lipid demixing. As neutral lipids are synthesized by enzymes located in the ER bilayer, they segregate from the phospholipids (PL) forming nanoscale lenses within the bilayer. With the subsequent incorporation of triglyceride molecules, these lenses grow into protuberances that eventually bud off from the ER membrane. Both lens formation and budding are key steps in LD biogenesis, and these processes have been successfully emulated by incorporating oil droplets into phospholipid vesicles (DEVs, for “Droplet Embedded Vesicles”) (1, 2) or by generating oil lenses in Langmuir monolayers (3), among other models. Using these systems, it has been possible to describe how LD formation and morphology are influenced by the interfacial tensions of both the bilayer and nascent LD (lenses). Highly curved lenses are more prone to bud off from the ER bilayer into independent LDs than flatter (less rounded) ones.

**S2. Rationale for choosing the Langmuir monolayer composition.** In a typical Langmuir monolayer experiment, a known amount of amphiphilic material dissolved in a volatile solvent insoluble in water, such as chloroform, is placed on the surface of water. After evaporation of the solvent, the monolayer material is compressed by mobile barriers that confine a variable interfacial area (A). The comparison between the surface tension at a given amphiphile density on the surface ($\gamma_{m}$) and the surface tension of the clean surface ($\gamma_{0}$) allow obtaining the lateral packing pressure $\pi$ as $\pi=\gamma_{0}-\gamma_{m}$. Knowing the number of molecules spread on the interface (n), allows estimating the Mean Molecular Area of the substances forming the monolayer phase, MMA=A/n. The laterally compression of monolayers allows studying π-MMA isotherms and characterizing molecular interactions, phase transitions, mixing or unmixing of components in the monolayer phase, among other properties. The "collapse" of the monolayer occurs at a certain pressure π_C_, where some molecules are no longer at the monolayer plane but forming 3D structures with internal structures depending on their composition, like folds, aggregates or lenses. In certain cases, one or more of the components of a film segregate into lenses, before the whole monolayer collapses; we will call π_EX_ to the value of π where this occurs. For instance, when the binary lipid system of PC/TG is used, TG lenses appear at π_EX_ as floating liquid structures with their interfacial tensions being in balance with that of the surrounding monolayer (see scheme below). There are various techniques to characterize Langmuir monolayers that provide diverse information, including Brewster Angle Microscopy (BAM). BAM allows the exploration of the organization of materials that make up the monolayers in two dimensions, including the size and shape of the domains and the heterogeneity of the Langmuir monolayers. It has been proved that the π_EX_ at which lenses are formed depends on PL/TG composition (4, 5). For instance, in the PC/TG mixture with a molar fraction of TG ($x_{TG}$) equal to 0.2 (PC/TG 4:1), lenses appear at πex 30-35 mN/m. Thus, in the present study we choose this composition as it corresponds to lateral packing comparable to those found in membrane bilayers.

**S3. The balance of interfacial tensions in the system of a floating lens in equilibrium with a surrounding film, as a model system representing a lens of neutral lipid inside a bilayer.** A lens embedded within a bilayer represents a nascent LD. In this system, the mechanical equilibrium arises from the balance of three interfacial tensions, each acting to minimize the area of its corresponding interface (Scheme 1, left panel). These are: (i) the interfacial tension acting at the bilayer ($\gamma_{b}$), and (ii) the two interfacial tensions acting at each lens–water interface ($\gamma_{LW}$). The interplay between these forces determines the shape of the lens following $\gamma_{b}=2\gamma_{LW}coscos \frac{\theta}{2}$ (according to the Neumann triangle of interfacial forces). For instance, when higher values of $\gamma_{b}$ will act by pulling the lens into a flatter shape, whereas lower $\gamma_{b}$values will favour the budding of a more rounded droplet into the aqueous phase.

The model system consisting of interfacial lenses in equilibria with a Langmuir film is a simpler physical basis for this model. It exhibits the same interfacial tensions equilibria (Scheme 1, right panel), but between the tensions present at ($\gamma_{m}$) the monolayer on the air/water interface, ($\gamma_{LA}$) the lens/air interface and ($\gamma_{LW}$) the lens/water interface. In a Langmuir film experiment, the monolayer tension is varied by lateral compressing the system while recording the interfacial tension or the lateral pressure π.

Although Langmuir films represent half a bilayer, they provide a valuable model to explore processes that occur in complete bilayers, with the advantage of enabling highly precise control over interfacial lateral packing (and tension) (3). In this framework, as π decreases in a Langmuir film that is in mechanical balance with lenses, this corresponds to an increase in $\gamma_{m}$ and thus, it induces lenses to flatten.

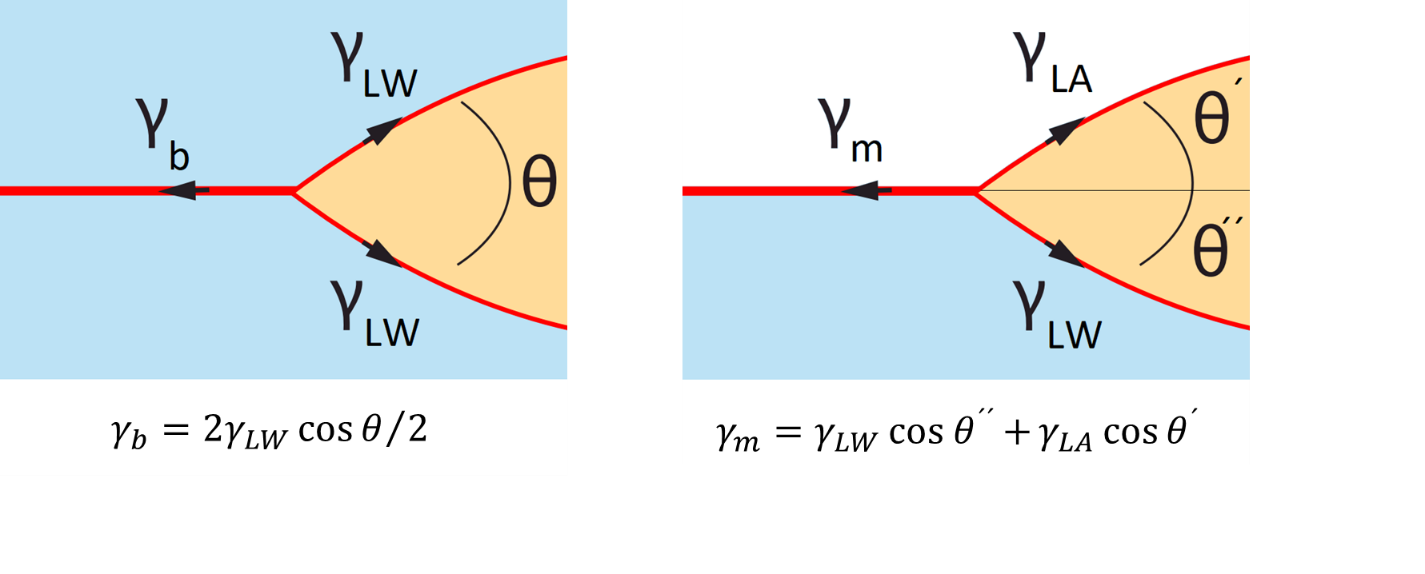

**Scheme 1:** Comparison of the three interfacial forces balancing each other on the equilibria of a lens with a surrounding bilayer or monolayer (in a Langmuir film). Adapted from (2).
